# A wireless, implantable optoelectrochemical probe for optogenetic stimulation and dopamine detection

**DOI:** 10.1101/2020.02.02.926782

**Authors:** Changbo Liu, Yu Zhao, Xue Cai, Yang Xie, Taoyi Wang, Dali Cheng, Lizhu Li, Rongfeng Li, Yuping Deng, He Ding, Guoqing Lv, Guanlei Zhao, Lei Liu, Guisheng Zou, Meixin Feng, Qian Sun, Lan Yin, Xing Sheng

**Affiliations:** School of Materials Science and Engineering, Beihang University, Beijing, 100191, China; Department of Electronic Engineering, Beijing National Research Center for Information Science and Technology and IDG/McGovern Institute for Brain Research, Tsinghua University, Beijing, 100084, China; Department of Physics, Tsinghua University, Beijing, 100084, China; Beijing Institute of Collaborative Innovation, Beijing, 100094, China; School of Materials Science and Engineering, Tsinghua University, Beijing, 100084 China; School of Optics and Photonics, Beijing Institute of Technology, Beijing, 100081, China; Department of Mechanical Engineering, Tsinghua University, Beijing, 100084, China; Key Laboratory of Nano-devices and Applications, Suzhou Institute of Nano-Tech and Nano-Bionics, Chinese Academy of Sciences (CAS), Suzhou 215123, China

## Abstract

Physical and chemical technologies have been continuously progressing advances of neuroscience research. The development of research tools for closed-loop control and monitoring neural activities in behaving animals is highly desirable. In this paper, we introduce a wirelessly operated, miniaturized microprobe system for optical interrogation and neurochemical sensing in the deep brain. Via epitaxial liftoff and transfer printing, microscale light emitting diodes (micro-LEDs) as light sources, and poly(3,4-ethylenedioxythiophene) polystyrene sulfonate (PEDOT:PSS) coated diamond films as electrochemical sensors are vertically assembled to form implantable optoelectrochemical probes, for real-time optogenetic stimulation and dopamine detection capabilities. A customized, lightweight circuit module is employed for untethered, remote signal control and data acquisition. Injected into the ventral tegmental area (VTA) of freely behaving mice, *in vivo* experiments clearly demonstrate the utilities of the multifunctional optoelectrochemical microprobe system for optogenetic interference of place preferences and detection of dopamine release. The presented options for material and device integrations provide a practical route to simultaneous optical control and electrochemical sensing of complex nervous systems.

## INTRODUCTION

In the past decades, the study of modern neuroscience has been significantly advanced, along with the monumental progresses of interdisciplinary technological innovations in fields including electrophysiology^1^, optical interrogation/detection^2^ and neuropharmacology^3^. Although traditional electrical tools for neural recording and medication provide powerful capabilities for brain science research and clinical therapies of neurological disorders like Parkinson’s disease and epilepsy^4^, the lack of cell-type specificity and side effects of stimulating untargeted brain regions impede their broader applications^5^. Recently developed genetically encoded optical actuators and indicators overcome the above limitations and offer more precise and cell specific neural signal regulation and monitoring^6, 7^. Additionally, multifunctional optical neural interfaces, comprising highly integrated, implantable microscale optoelectronic and microfluidic devices, have achieved remarkable accomplishments in optical/electrical/chemical multimodal sensing and modulation in the deep brain^8–12^. Representative examples include our recently developed optoelectronic probes for wireless optogenetic stimulation and fluorescence recording in freely moving rodents^13–17^. Besides electrical pulses and calcium flows, neurotransmitters, which comprise a plethora of chemicals including dopamine, glutamate, serotonin, etc., are of critical importance and direct relevance in neural activities and brain functions^18, 19^. Among the family of neurotransmitters, dopamine arouses particular interests because of its tight association with important behaviors like motivation, reward, and reinforcement^20^. For example, morphological and neurochemical studies have noted prominent reductions of dopamine in the striatum of patients with Parkinson’s disease, and identifying the relationship between dopamine release and neuron stimulation is crucial for understanding the disease cause and developing precise treatments.^21, 22^ Therefore, it is highly desirable to develop a fully integrated implantable system leveraging optogenetic stimulators and dopamine sensors for closed-loop neural stimulation and recording. A commonly used method for dopamine detection involves tethered electrochemical probes made of carbon or noble metals like gold and platinum, which could potentially be bundled with optical fibers for optical simulation^23–25^. More recently, genetically encoded fluorescence indicators of various neurotransmitters have also been actively exploited^26–29^. Integrating the detection of neurotransmitter levels within the wirelessly operated optogenetic stimulation platforms in living animals, however, is relatively underexplored in state-of-the-art studies.

In this paper, we develop a wireless thin-film based implantable microprobe system for optogenetic stimulation and electrochemical sensing of dopamine in the animal deep brain. A thin-film, microscale light-emitting diode (micro-LED) transferred on a flexible substrate is employed as a light source for optogenetic stimulation. A poly(3,4-ethylenedioxythiophene) polystyrene sulfonate (PEDOT:PSS) coated diamond film is placed upon the micro-LED, serving as an electrochemical sensor for dopamine detection. As an optically transparent, electrically insulating and thermally conductive interlayer in the stacked device structure, the diamond film separates the micro-LED and the PEDOT:PSS sensor, ensuring simultaneous optical stimulation and electrochemical detection in the same region. For wireless control and signal acquisition, the formed thin-film and flexible microprobe is interfaced with a miniaturized, removable circuit module powered by a rechargeable battery. Implanted into the ventral tegmental area (VTA) of freely behaving mice, the fully integrated optoelectrochemical microprobe is demonstrated to remotely control animal behaviors and dopamine signal recording.

## RESULTS AND DISCUSSION

### Device structure, fabrication and functionality

Fig. 1a schematically displays the conceptual illustration of our proposed implantable, fully integrated microprobe system for simultaneous optogenetic stimulation and electrochemical sensing in the deep brain tissue, with an exploded view shown in Fig. 1b. Details about the device structure and fabrication processes are provided in the supplemental information (device fabrication, and Figs. S1–S3). The thin-film device stack incorporates a flexible double side copper (Cu) coated polyimide (PI) (18 μm Cu / 25 μm PI / 18 μm Cu) substrate, an indium gallium nitride (InGaN) based blue emitting micro-LED (size: 125 μm × 185 μm × 7 μm), an undoped diamond interlayer (size: 180 μm × 240 μm × 20 μm) and a PEDOT:PSS film (size: 150 μm × 200 μm × 0.1 μm). Here we employ the side Cu coated PI film instead of the pure PI as the supporting substrate, because of the desirable heat dissipation of Cu films (Fig. S4). The emission from the blue LED is utilized to optically control specific cells expressing light-sensitive ion channels, for example, channelrhodopsin-2 (ChR2) ^6, 7, 30–32^, thereby activating or inhibiting neural activities. As an optically transparent and electrically conductive polymer layer, the PEDOT:PSS thin film can work as an effective electrochemical sensor for dopamine detection, owing to its biocompatibility and low impedance at the biological interface^33–36^. In Fig. 1b, the PEDOT:PSS film serves the working electrode (WE), while the reference electrode (RE) and the counter electrode (RE) are respectively based on a standard Ag/AgCl wire and a platinum wire, which are integrated separately to reduce the device footprint. The optical transmittance of the PEDOT:PSS film is measured to be more than 90% in the visible spectral range (Fig. S5). As an interlayer between the LED and the PEDOT:PSS sensor, the benefits of using the diamond film are manifold: 1) Its optical transparency (> 80%)^37^ ensures that the blue light can transmit through it and reach the bio-tissue; 2) Its high thermal conductivity (> 2000 W/m/K)^38^ allows for efficient heat dissipation during LED operation; 3) It electrically isolates the electrodes of the LED and the PEDOT:PSS film, minimizing signal crosstalk; and 4) It could act as a liquid barrier and prevent the LED from failure in the aqueous environment. In addition, we note that the lateral dimensions of the diamond film are designed to be slightly larger than the micro-LED, in order to: 1) facilitate heat dissipation; 2) provide better encapsulation on the LED; and 3) improve the misalignment tolerance during the transfer and lithography processes. The LED is fabricated on sapphire and the diamond film is grown on silicon. They are released from native substrates respectively by laser liftoff and wet etching, forming freestanding films transferred onto PI using polydimethylsiloxane (PDMS) stamping methods^39, 40^. On the diamond layer, the PEDOT:PSS film is conformally coated via spin casting and lithographically patterned, with a thickness of ∼100 nm (Fig. 1c). The micro-LED and the PEDOT:PSS film are metalized with sputter coated Cr/Cu/Au and Cr/Au electrodes respectively, while epoxy-based photoresist films (SU-8) serve as the bonding layer between the LED and the diamond, as well as the protective coating on metal electrodes. Finally, the flexible substrate is milled by ultraviolet laser to realize a needle shape (width ∼ 360 μm, length ∼ 5 mm, shown in Fig. 1d), which is ready for implantation into the deep tissue of animal brains. Compared with other possible device geometries like physically bonded fibers/wires^41, 42^ or laterally placed thin-film sensors^16, 43, 44^, the vertically stacked device structure is more advantageous because: 1) It has a higher spatial precision in the control and the monitoring of a specifically targeted nucleus, which typically have sizes from tens to hundreds of micrometers in the mouse brain; and 2) Such a thin-film geometry can have a smaller footprint and desirable mechanical flexibility^45^, significantly reducing tissue lesion and associated damages.

**Figure 1.**
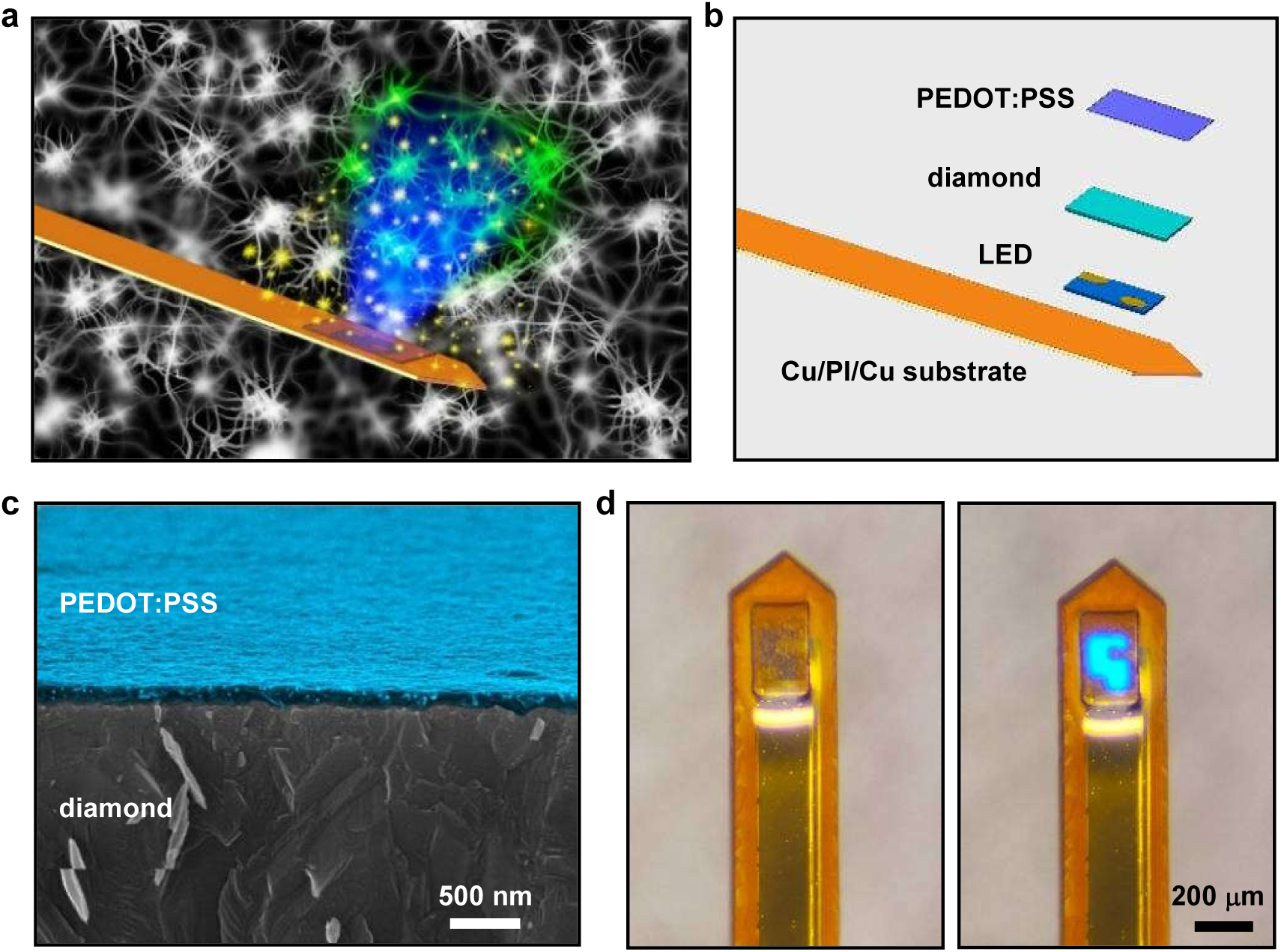
Structures of the implantable microprobe. **(a)** Conceptual illustration of the microprobe system for optogenetic stimulation and dopamine detection. **(b)** Exploded schematic of the microprobe, comprising a blue micro-LED, a diamond interlayer and a PEDOT:PSS thin film on the copper coated polyimide (Cu/PI/Cu) substrate. **(c)** Cross-sectional scanning electron microscope (SEM) image of a PEDOT:PSS film (colorized in blue) coated on diamond. **(d)** Optical photographs of a fully fabricated microprobe (left: LED is off; right: LED is on, with an injection current of 0.1 mA).

### Optoelectronic and thermal properties

The utility of freestanding InGaN based micro-LEDs for optogenetic stimulations has been demonstrated in previous works^40^. Here we evaluate optical and thermal properties of the micro-LED before and after integrating with the PEDOT:PSS coated diamond film for electrochemical sensing, with results depicted in Fig. 2. Under current injection, the peak wavelength for the LED emission is around 470 nm (Fig. 2a), which matches to the optical absorption of commonly used opsins like ChR2 as optogenetic tools^15, 30–32^. The external quantum efficiency (EQE) for LEDs with and without diamond and PEDOT:PSS films are plotted in Fig. 2b. Maximum EQEs are ∼12% (for the bare LED) and ∼10% (for the LED coated with diamond and PEDOT:PSS), obtained at an injection current of ∼1 mA. In addition, both devices exhibit nearly Lambertion emission distributions (Fig. 2c). These results indicate that the diamond/PEDOT:PSS film placed on top of the blue LED does not greatly alter its emission characteristics, and the slight performance degradation (∼20%) is in accordance with the high optical transmission of the diamond and the PEDOT:PSS materials (>80%)^37, 46^.

**Figure 2.**
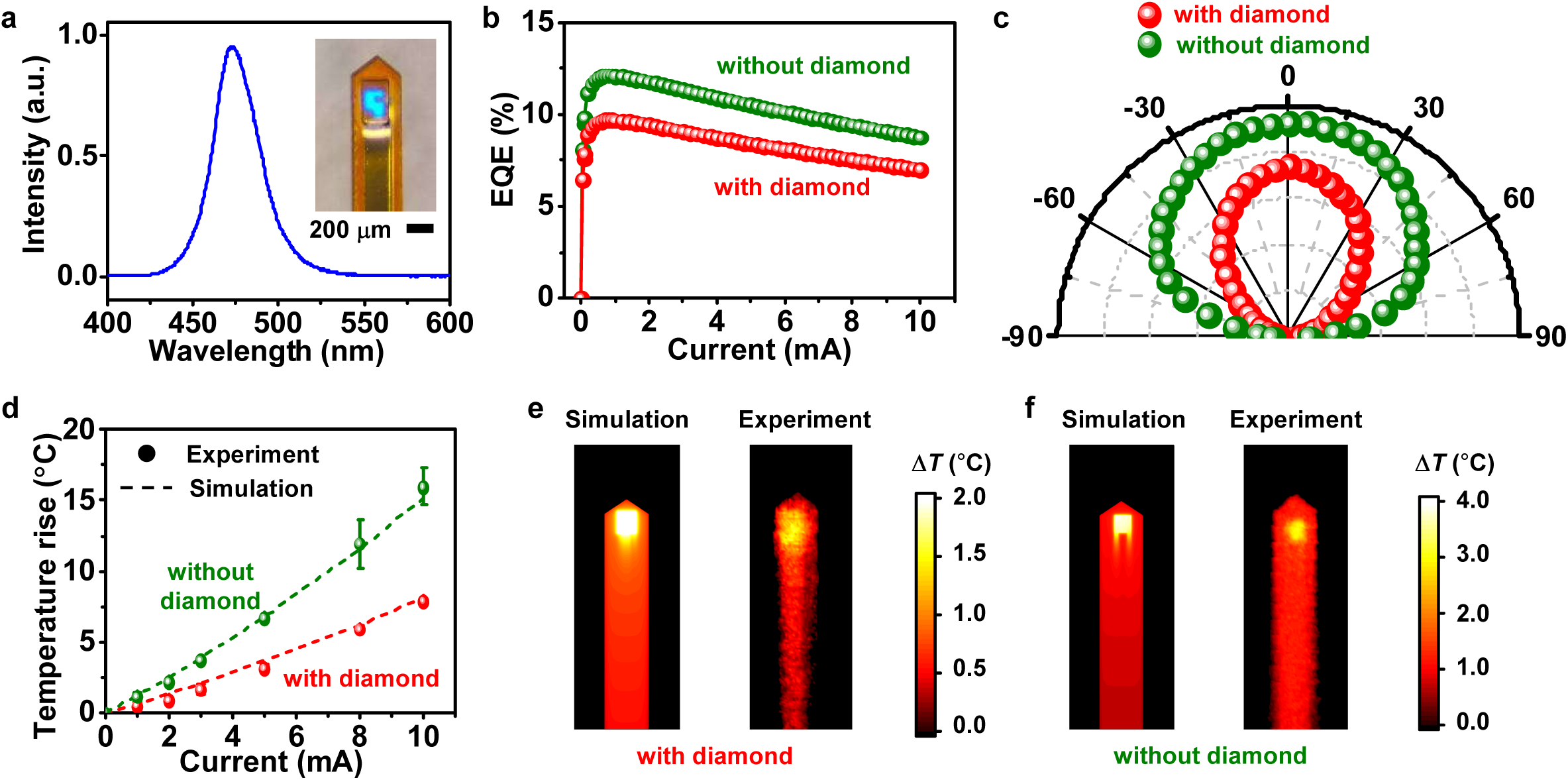
Optoelectronic and thermal characteritics of LEDs in the microprobe. **(a)** Electroluminescence spectrum of the InGaN blue LED. Inset: optical photograph of the microprobe. **(b)** External quantum efficiencies (EQEs) for LEDs with and without diamond/PEDOT:PSS coatings, under different injection currents. **(c)** Angular distribution of optical emissions for LEDs with and without diamond/PEDOT:PSS coatings. **(d)** Measured (dots) and simulated (dashed lines) maximum temperature rises above room temperature on the top surface of the LED probes (with and without diamond/PEDOT:PSS coatings) as a function of pulsed currents (frequency 20 Hz and duty cycle 20%). **(e, f)** Simulated (left) and measured (right) temperature distributions the LED probes (current 3 mA, frequency 20 Hz and duty cycle 20%), **(e)** with and **(f)** without diamond/PEDOT:PSS coatings.

Besides optical transparency, another prominent feature of the diamond film is its ultrahigh thermal conductivity. Even through the diamond film we apply here has a polycrystalline morphology, its thermal conductivity still reaches as high as 2200 W/m/K, almost 5 times higher than copper films and 3 orders of magnitude higher than organic coatings like SU-8 and PDMS^47^. Such a thermally conductive coating is advantageous for heat dissipation of LEDs operated in biological systems. In Fig. 2d–f, we evaluate the thermal behaviors for LEDs with and without diamond/PEDOT:PSS coatings. The LEDs are operated in ambient air under pulsed current injection (varied from 0 to 10 mA), with 20 Hz frequency and 20% duty cycle. Temperature maps are measured with an infrared camera and compared with thermal modeling established by finite element analysis. As shown in Fig. 2d, the maximum temperature rises for the LED with diamond are approximately one half of those for the bared LED under the same injection currents. Thermal mapping results for LEDs operated at 3 mA are further depicted in Fig. 2e and 2f. These experimental results are in good agreement with numerical predictions. At 3 mA, the probe surface temperature increase is restricted within 2 °C for the LED with the diamond coating, while the result for the bare LED is more than 4 °C. Furthermore, it should be noted that for the microprobe implanted in biological tissues, the temperature rises, albeit more challenging to measure precisely, should be even lower than the results obtained in air, because of the additional heat dissipation capability of biological tissues and fluids. The simulation results based on the established models are provided in Fig. S6. This benefit is particularly favorable for optogenetic probes implanted in the deep brain, since the mitigation of unwanted abnormal activities and possible tissue damaging by overheating is critical for neurons^48^.

### Electrochemical characterization

Electrochemical properties of the PEDOT:PSS electrode for dopamine sensing are studied and presented in Fig. 3. Although the normal dopamine level in the whole animal brain or other body parts is as low as a few pM^49, 50^, it can be much higher (from several nM to several μM) in particular brain regions with large collections of dopaminergic neurons^51, 52^. The detection of dopamine in aqueous solutions and biological environments is based on its redox reaction, in which dopamine is oxidized and converted into dopamine-o-quinone on the electrode surface, generating electric currents^53^. Cyclic voltammetry (CV), differential pulse voltammetry (DPV), chronoamperometry (CA), and fast scan cyclic voltammetry (FSCV) techniques are utilized to analyze the electrochemical features of PEDOT:PSS electrode, using a standard electrochemical work station. A standard three-electrode configuration is applied, with the PEDOT:PSS film serving as the working electrode (WE), silver/silver chloride (Ag/AgCl) and platinum (Pt) as the reference (RE) and the counter electrodes (CE), respectively. Here we use dilute hydrochloric acid (HCl) solutions (pH = 4.0) as solvents for dopamine, in order to mitigate its degradation due to natural oxidation in air that causes measurement inaccuracy^54, 55^. The dopamine concentrations used in CV tests are 0 μM, 0.5 μM, 1 μM, 10 μM, 50 μM, 100 μM and 500 μM. As shown in Fig. 3a, pairs of well-defined quasi-reversible redox peaks are observed in the presence of dopamine in HCl, with oxidation potentials of 0.5–0.6 V and the corresponding reduction potentials of 0.3–0.4 V. Peak redox currents increase with the concentration of dopamine in the solution. DPV measurements in Fig. 3b reveal similar trends, with a higher sensitivity of concentration and a better voltage resolution than those obtained with CV analysis. Dopamine levels in HCl solutions as low as ∼0.1 μM can be detected using DPV, and the detection limit using CV and CA are ∼0.5 μM. Repetitive CV scanning curves of a representative probe are shown in Fig. S7, demonstrating the reliability and repeatability of its electrochemical properties. DPV curves for multiple probes are also presented (Fig. S8), in which the response differences among various probes are ascribed to the variations in the fabrication process (spin coating, misalignments in lithography and etching, etc.), as well as the instability of dopamine in the ambient environment. The dynamic current response at varied dopamine concentrations is further determined using CA measurements in Fig. 3c and 3d. At a fixed bias voltage of 0.6 V, oxidation currents are captured as we change dopamine concentrations. In the range of 0.1–10 μM, a linear relationship between the dopamine concentration and the response current can be obtained, and the detection sensitivity is determined to be ∼0.06 nA/μM. For our spin coated PEDOT:PSS film with a working area of 150 μm × 200 μm defined by lithography, the normalized current response of dopamine by area is calculated to be 200 nA/μM/cm^2^, which is comparable to that measured by electrochemical sensors based on other materials^56, 57^. In Fig. 3e, we further record the FSCV responses by dropping dopamine solutions in HCl, plotting the current response as a function of both time and voltage. It should be noted that CV, DPV, CA and FSCV measurements exhibit different current sensitvities at varied dopamine concentrations, because part of the measured electrochemical currents are associated with the capacity at the electrode/electrolyte interface, which is highly sensentive to the applied applying different potential modulation forms and scan rates^58^. The experimental results performed *in vitro* clearly demonstrate the utility of applying the PEDOT:PSS film for real-time dopamine monitoring in biological systems.

**Figure 3.**
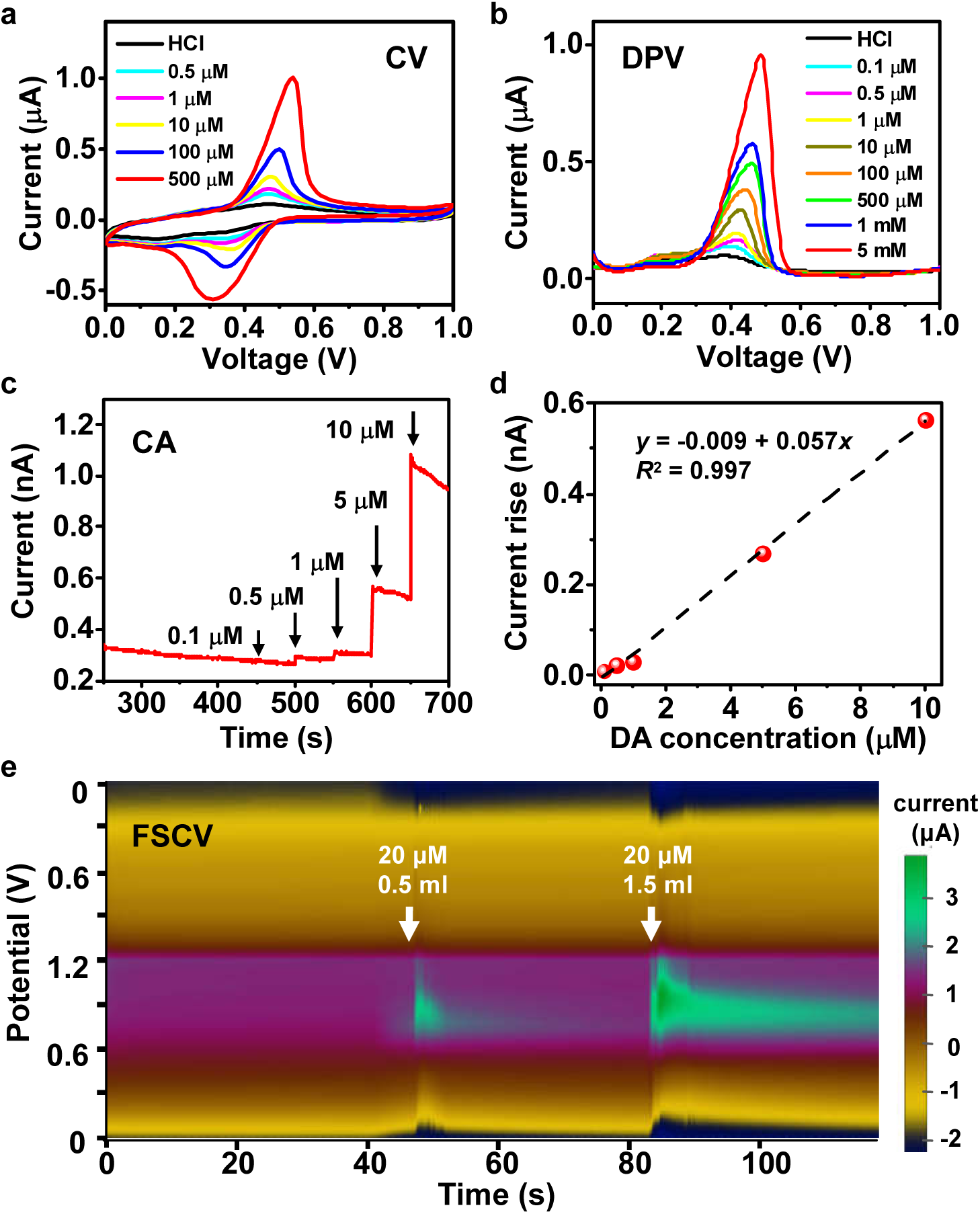
Electrochemical characteristics of the PEDOT:PSS film in the microprobe. **(a)** Cyclic voltammetry (CV). **(b)** Differential pulse voltammetry (DPV). **(c)** Chronoamperometry (CA). **(d)** Calibration curve: linear relationship between response currents and dopamine (DA) concentrations (0.1–10 μM). **(e)** Fast scan cyclic voltammetry (FSCV). All the experiments are performed in HCl solutions (pH = 4.0) with varied dopamine concentrations at room temperature, with a saturated Ag/AgCl electrode as the reference and a Pt sheet as the counter electrode.

### Circuit design and evaluation

A customized circuit module is designed to wirelessly operate the implantable microprobe for integrated optoelectrochemical stimulation and sensing functionalities. The circuit diagram is depicted in Fig. 4a, with detailed layout presented in Fig. S9 and S10. Remote data communication is realized with a micro controller (nRF24LE1) with a transceiver operated at 2.4 GHz. A driver chip (ZLED7012) provides a constant current to the micro-LED for optogenetic stimulation, with programmable current levels, frequencies, pulse widths, etc. A red LED is printed on the circuit outside the brain for signal indication. A digital-analog converter (DAC60508) and two pre-amplifiers (OPA2381 and ADA4505) are connected to the chemical electrodes (WE, RE and CE) for voltage scanning and current readout, achieving a resolution as low as 0.1 nA for current measurements. The analog signals from the electrodes are collected, filtered, compared and amplified in the operational network and send to the corresponding ports of the micro controller. After further data process via the built-in program, the signals are wirelessly transmitted to the receiving end connected to a laptop computer via an antenna operated at a radio frequency of 2.4 GHz (Fig. S11). The system is powered by a rechargeable lithium ion battery with a capacity of 35 mAh. As shown in Fig. 4b, the microprobe consisting of a blue micro-LED and a PEDOT:PSS coated diamond film as an electrochemical sensor is connected to the wireless circuit module with a footprint of ∼2.2 cm × 1.3 cm and a weight of 2.0 g (including a 0.9-g battery). Via a flexible printed circuit connector, the manufactured circuit can be plugged into the implantable microprobe during *in vivo* experiments and detached from it when unnecessary. We evaluate the performance of our designed wireless circuit *in vitro* by taking CV scans in HCl solutions with various dopamine concentrations, using a microprobe containing the PEDOT:PSS thin-film electrode. The measured CV results in Fig. 4c match reasonably well with those obtained using a commercial electrochemical analyzer, with clearly resolved redox peaks of dopamine. Similarly, CA measurements (current versus time at a fixed bias voltage) can also be performed. These results demonstrate the utility of the wireless circuit for *in vivo* tests in behaving animals.

**Figure 4.**
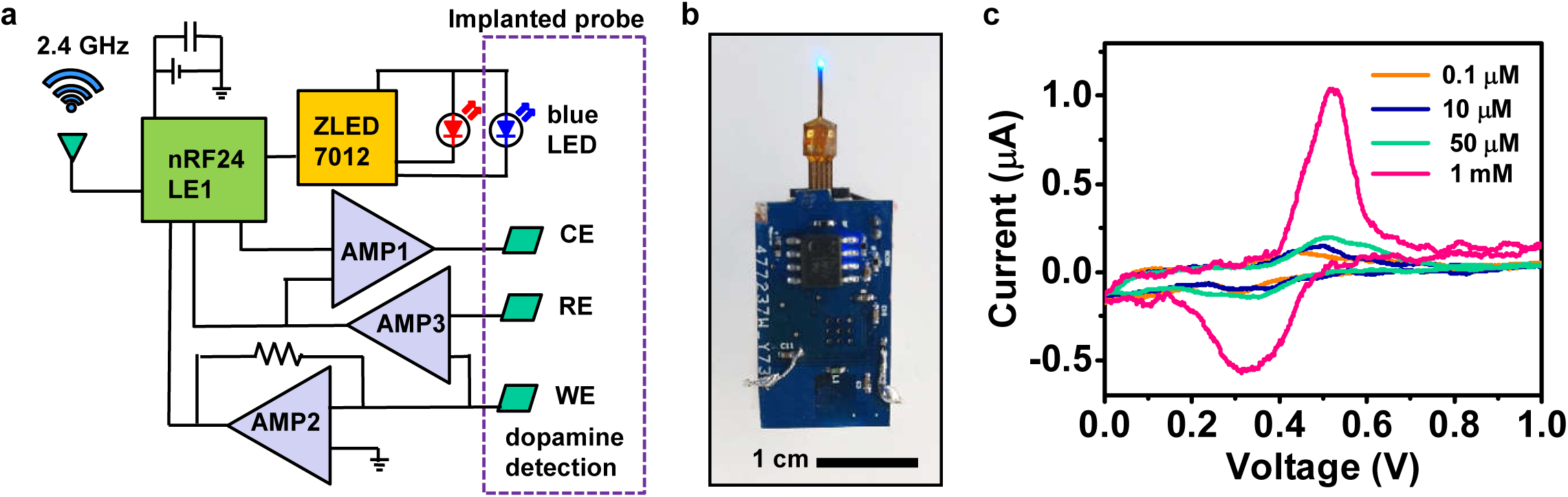
Circuit module for wireless operation. (a) Schematic diagram of the circuit. (b) Photograph of a circuit connected to an implantable probe with the micro-LED on. (c) CV curves acquired by the wireless circuit connect to a microprobe in HCl solutions with different dopamine levels.

### *In vivo* experiments

We further verify the performance of the wirelessly operated microprobe with *in vivo* optogenetic stimulation and dopamine sensing in experimental mouse models. The system can be easily mounted on the head of an adult mouse, without affecting its behaviors (Fig. 5a). Illustrated in Fig. 5b, probes including micro-LEDs and PEDOT:PSS electrodes are implanted into the ventral tegmental area (VTA), which is the origin of dopaminergic cells and involves reward circuitry^59, 60^. Previous studies show that dopamine concentrations in the VTA typically range from several nM to several μM, depending on local neural acitivities^51, 52^. A microprobe with a length of ∼5 mm is implanted into the VTA of adult mice (DAT-Cre transgenic, 7–16 weeks) expressing ChR2, with detailed surgery procedures provided in Figs. S12 and S13. Fig. 5b shows a brain section with the lesion area created by the implanted microprobe. Details of the immunostained region in the VTA are provided in Fig. 5c, including stained ChR2 (red), rabbit polyclonal anti-tyrosine hydroxylase (TH, green), 4’,6-diamidino-2-phenylindole (DAPI, blue) and merged images. The track of the probe is indicated by the dashed boxes. Tissue damages and inflammatory responses are similar to those induced by the insertion of microprobes with similar geometries and flexible materials reported previously^16, 61^. We perform real-time place preference (RTPP) tests by optogenetic stimulation 5 days after implanting microprobes into the VTA of mice. Mice are placed in a behavior box (50 cm × 25 cm × 25 cm) with two chambers and a gate with a width of 9 cm. Animals are allowed to explore the environment for 15 minutes to get a pretest result, and only those with no obvious place preference (Fig. 5d, upper part) are used for optogenetic test. Recorded position heat maps and traces (Fig. 5d, as well as more raw data provided in Fig. S14) clearly show that the implanted microprobe and the mounted wireless module impose few constraints on the animals’ locomotion. After pretest, optical stimulation (duration: 10 ms, frequency: 20 Hz, driving current: 1.8 mA) generated by the blue-emitting micro-LED is provided for 15 minutes when mice go into the left chamber. The position heatmap (Fig. 5d, lower part) reveals that the mice exhibit obvious place preferences during the optogenetic stimulation period. Statistical results clearly show that animals spend more time in the stimulated chamber and have obvious place preference compared to pretests, indicating that dopaminergic neurons in the VTA are excited and dopamine release is promoted. The optogenetic stimulation can be effective more than two weeks after the micro-LED implantation (Fig. S15). In fact, the implanted micro-LEDs and ChR2 expression can be stable for months after surgery, as previously demonstrated^14^.

**Figure 5.**
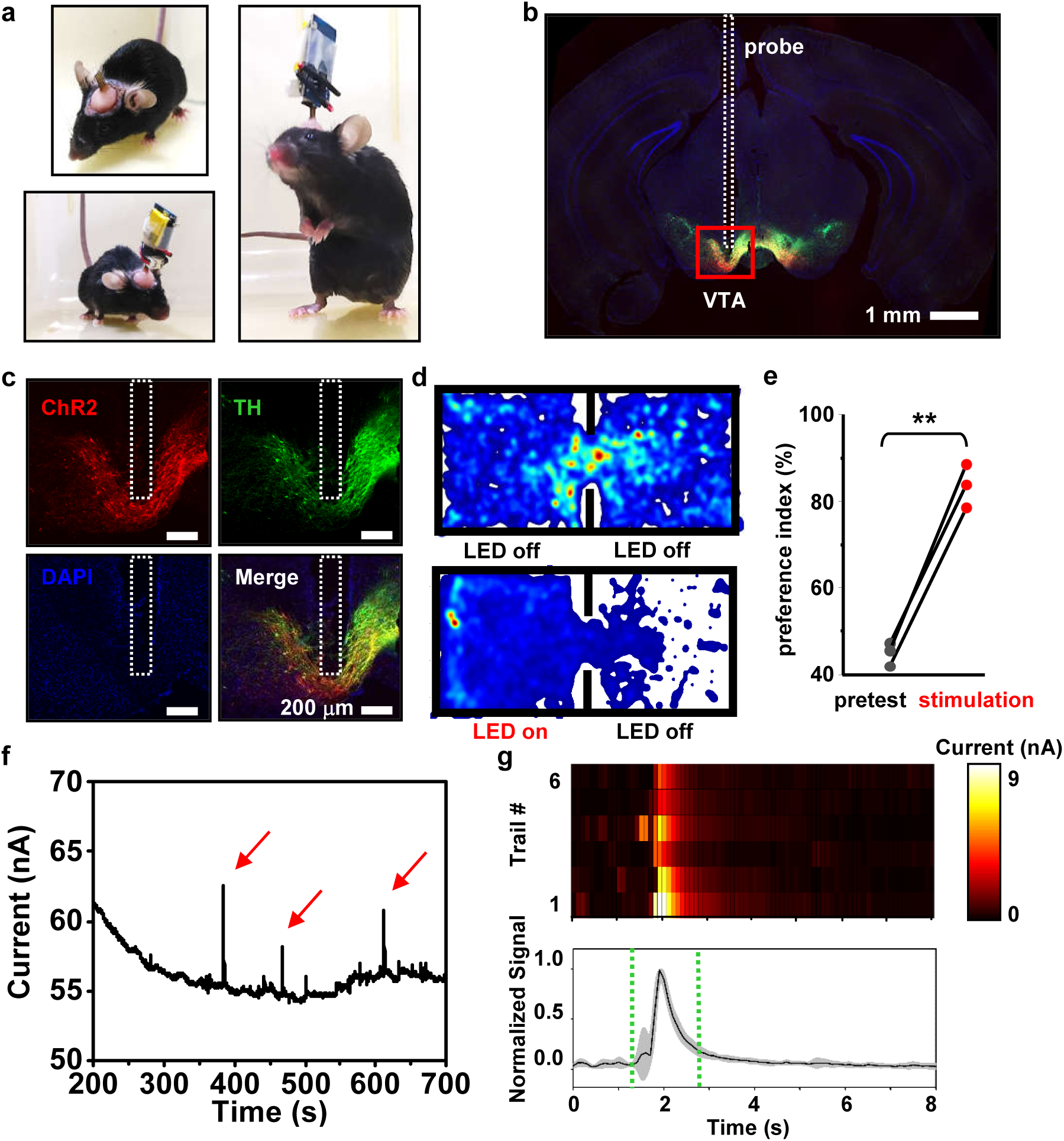
*In vivo* optogenetic stimulation and dopamine detection. (a) Photographs of an experimental mouse implanted with the microprobe, with and without the circuit module. (b) Immunostained fluorescence image of the brain section. (c) Immunostaining results of the VTA region expressing ChR2 (red), TH (green), DAPI (blue), as well as the merged image. (d) Position heat maps of the animal activity during pretests (top) and under optogenetic stimulations (bottom). Hotter colors represent longer duration in the location. (e) Preference indices (the ratio of the time that mice spend in the left chamber to the whole recorded time) on the left side under optogenetic stimulations, in comparison with those during pretests (*n* = 3 mice, unpaired *t* test, *P* < 0.01). Stimulation parameters: duration 10 ms, frequency 20 Hz, driving current 1.8 mA). (f) A representative amperometry curve (current versus time) measured with the PEDOT:PSS electrode in vivo. (g) Analyses of representative signal peaks. Top: Heatmaps showing 6 individual trials (current signals vs. time) from 2 mice. Bottom: Average of normalized current signals.

To evaluate the electrochemical sensing capability, we measure CA curves (current versus time) with the PEDOT:PSS electrode implanted in the VTA. The VTA contains a mixture of dopaminergic (∼65%), GABAergic (∼30%), and glutamatergic (∼5%) neurons^62^. Both Gamma aminobutyric acid (GABA) and glutamate are non-electroactive^63, 64^, thus making dopamine the dominate neurotransmitter probed in electrochemical tests in the VTA. Similar to *in vitro* experiments in Fig. 3, here we employ a Ag/AgCl wire and a stainless steel screw respectively as the reference and counter electrodes, which are implanted elsewhere in the mouse brain cortex. Here the stainless steel screw is a standard fixture for the implantation surgery, so we directly use it as the counter electrode so that we do not have to implant another platinum wire, which causes additional tissue damage. A constant forward bias of 0.6 V is applied between the WE and the RE, and the oxidation current flowing into the WE is acquired at an interval of 0.1 s. Fig. 5f plots a representative amperometric results measured with a microprobe implanted into the VTA of mice. Spontaneous current spikes are observed, and more signal trails from multiple animals are collected and analyzed in Fig. 5g (with raw data shown in Fig. S16). The data collected from multiple individual trials reveal considerable consistency, with time constants matched to dopamine signals reported in other literatures^65–67^. Here we also observe that the micro-probe can only be properly functional under acute tests *in vivo*, and its electrochemical sensitivity gradually degrades several hours after implantation, because of potential chemicals and bio-substances adhering to the probe surface (as an example shown in Fig. S17). Similar electrode fouling effects have also been reported for other electrochemical sensors based on, for example, carbon fibers^68^. As a future research thrust, this challenge needs to be addressed to enable chronic electrochemical detection *in vivo*.

## Conclusion

In summary, we report a wirelessly operated microprobe system for neural interrogation and neurotransmitter monitoring in the deep brain. Ultraminiaturized, vertically stacked micro-LED, diamond and PEDOT:PSS films are combined to realize optogenetic stimulation and dopamine sensing in a minimally invasive platform. The unique electrical, optical and thermal properties of the PEDOT:PSS coated diamond film make the device suitable for highly sensitive electrochemical sensing, without affecting the micro-LED operation. A lightweight, remotely control circuit allows for behavior studies on freely moving mice. In the future, more sophisticated *in vivo* experiments can be performed to demonstrate closed-loop operations, for example, modulate light stimulation in response to the change of dopamine levels, or monitor the dopamine release when varying optical emissions. Endogenous dopamine release associated with various reward behaviors can also be probed. Moreover, the device capability can be further evaluated in animals with relevant disease models such as Parkinson’s disease and depression^61, 69^, in which optogenetic stimulation can be utilized to activate or inhibit dopaminergic neurons to precisely control dopamine release in the brain. Advanced microfluidic device can also be integrated into the platform for automated drug delivery, enabling pharmacological modulation of neural activities^11, 12^. It should be noted that the sensing capability of the PEDOT:PSS electrodes may be interfered by the micro-LED operation due to possible optoelectronic artifacts similar to those reported on other optotrodes^67, 70^. In fact, we have observed this kind of signal interferences in experiments (Fig. S18). To resolve these issues, strategies including modifying circuit design and incorporating new signal processing algorithms^71, 72^ will be investigated in the future. Additionally, multiple microprobes can be implanted into different brain regions in one or multiple animals, exploring function connectivities among distinct units within the nervous system. The sensitivity to the dopamine concentration can be further optimized, by adjusting the properties of the PEDOT:PSS film, including its geometries, doping, surface treatment, etc. Considering that various neurotransmitters such as glutamate, serotonin and norepinephrine, existing in the neural system, specificity and sensitivity of dopamine can be further improved by selective coatings including metal nanoparticles (Au^73, 74^, Pt^75^, Pd^76^), enzyme (laccase^77^, tyrosinase^78, 79^) and carbon-based material^80, 81^ on the PEDOT:PSS^79^. As for most electrochemical sensors used for *in vivo* experiments, trade-offs between sensitivity, selectivity and stability should be taken into account^82, 83^. Although here we only demonstrate acute tests in living mice, chronic operation of electrochemical sensors is also a key challenge for complex behavior experiments. Besides optogenetics and electrochemistry, many research tools including fluorescence photometry^16^, electrophysiology^84^, and microfluidics^85^, can be combined in such a platform, to accomplish multimodal electrical-optical-chemical closed-loop modulation and detection of neural activities. Additionally, the circuit design can be further optimized, for example, by incorporating flexible circuit boards and battery-free energy harvesting techniques^85–87^. Collectively, we envision that these advanced strategies provide a promising route to tackle unresolved challenges in fundamental neuroscience and medical practices.

## MATERIALS AND METHODS

### Device fabrication

Details about device structures and fabrication processes are provided in the supplemental information (device fabrication, and Figs. S1–S3).

Micro-LEDs: Details for the micro-LED fabrication are described in the previous work^40^. The InGaN-based blue LEDs are epitaxial grown on sapphire substrates using metal-organic chemical vapor deposition (MOCVD) and lithographically formed. The structure (from bottom to top) consisted of a GaN buffer layer, an n-GaN layer, an InGaN/GaN multiple-quantum-well layer, and a p-GaN layer, with a total thickness of about 7.1 μm. The lateral dimension of LED is defined by inductively coupled plasma reactive ion etching (ICP-RIE), with 4 mTorr pressure, 40 sccm Cl_2_, 5 sccm BCl_3_, 5 sccm Ar, ICP power 450 W, bias power 75 W, and an etch rate of 0.33 μm/min. Freestanding thin-film micro-LEDs (dimension: 180 μm × 125 μm × 7 μm) are formed by laser liftoff (Light Source: the KrF excimer laser at 248 nm, Coherent, Inc., CompexPro110). The optimized power density for the laser is around 0.7 J/cm^2^.

Freestanding diamond film: The polycrystalline diamond film (thickness ∼ 20 μm) is grown on single crystalline silicon substrates via chemical vapor deposition (CVD). Laser milling (Nd:YVO4 laser, 1064 nm) is used to cut the film into designed patterns (size: 180 μm × 240 μm) and shapes. The silicon substrate is removed using an etchant solution (CH_3_COOH : HNO_3_ : HF = 5 : 5 : 2 by volume ratio), realizing freestanding diamond films.

Fabrication of the microprobe: A 10-µm thick polyimide is coated on the Cu/PI/Cu substrate and baked at 250 °C for over 120 mins as an insulating layer between the micro-LED and the substrate. Pre-patterned polydimethylsiloxane (PDMS) (Dow Corning Sylgard 184 kit, 1:10 weight ratio) stamps, the freestanding LEDs are transferred onto flexible double side Cu coated polyimide (18 μm Cu / 25 μm PI / 18 μm Cu, from DuPont) substrates, with a spin-coated adhesive layer. LEDs are encapsulated with 5 μm thick SU-8 (SU8-3005) as an insulating layer. Sputtered layers of 10 nm Cr / 500 nm Cu / 100 nm Au serve as the metalized electrode for the micro-LEDs, and coating 5 μm thick SU-8 (SU8-3005) as an encapsulation layer. The freestanding diamonds are transferred onto micro-LEDs using similar PDMS stamps with 2 μm thick SU-8 (SU8-2002) as an adhesive layer. A 5 μm thick SU-8 (SU8-3005) film is spin coated on the sample, with the surface of the diamond lithographically exposed. Sputtered 5 nm Cr / 500 nm Au films are used as interconnected electrodes for the PEDOT:PSS films. The PEDOT:PSS film is coated on the diamond as the working electrochemical electrode, and patterned by reactive ion etching (RIE) (90 mTorr pressure, 100 sccm O_2_, 5 sccm SF_6_, power 100 W for 200 s). Another photoresist film (SU8-3005, 5μm thick) is patterned on the metal electrodes as a waterproof layer. Subsequently, the flexible substrates are patterned to form needle shapes (width ∼360 μm, length ∼ 5 mm) by UV laser milling.

### Device characterization

SEM images are captured by ZEISS Merlin microscope (15 kV). Optical microscopy images are taken by microscope MC-D800U(C). The EQE of micro-LEDs are measured using an integrating sphere (Labsphere Inc.) and a calibrated Si photodetector. Thermal images are acquired by an infrared camera (FOTRIC 228). LED emission spectra are collected with a spectrometer (Ocean Optics HR2000+). Far-field angular emission profiles of micro-LEDs are captured by a standard Si photodetector (DET36A, Thorlabs). *In vitro* electrochemical tests of dopamine detection are performed with an electrochemical work station (CHI 650E, Shanghai Chenhua Co., Ltd, China). The CV is performed at a sweep rate of 0.1 V/s from 0 V to 1 V. Conditions for the DPV are, voltage step: 4 mV, pulse time: 200 ms, pulse amplitude: 50 mV, and scan rate: 20 mV/s. The CA is employed at a constant bias of 0.6 V. The FSCV is performed with a sweep rate of 10 V/s from 0 V to 1.2 V for 2000 circles. All of the electrochemical measurements are carried out in a Faraday cage to avoid electromagnetic disturbance from environments.

#### Thermal Simulation

The simulations of temperature increase on the surface of LED probes (with and without diamond) are performed by steady-state finite element methods in COMSOL Multiphysics®, using the module of Heat Transfer in Solids. The three-dimensional geometric model of the probed is constructed based on the structure described in this paper. The InGaN LED is defined as the heat source, with the input thermal power density (W/m^3^) estimated by

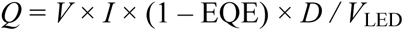

where *V* and *I* are measured voltage and current of the LED under continuous current injection conditions, EQE is the external quantum efficiency, *D* is the duty cycle of LED in experiments (20% in this study) and *V*_LED_ is the volume of the LED. Boundary conditions of natural convection to air are applied to all external surfaces of the probe model, in accordance with the experiment setup, and the air temperature is set to 20 °C. The temperature distribution T is calculated by solving the following heat transfer equation:

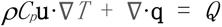

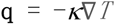

where *ρ* is the material mass density, *C_p_* is the heat capacity at constant pressure and *κ* is the thermal conductivity. A self-adaptive mesh setup is applied on the geometric model. Solutions of temperature are acquired by a stationary fully-coupled solver within the convergence tolerance of 10^-5^ using Newton iteration methods, and the final results of temperature changes are plotted on the external surface of the probe model from top view.

### *In vivo* studies

#### Stereotaxic surgery

Animal protocols are approved by Animal Care and Use Committee of Tsinghua University and the National Institute of Biological Sciences, Beijing. Adult (7∼16 weeks) DAT-Cre transgenic mice (Jackson Laboratory, strain B6.SJL-Slc6a3tm1.1(cre)Bkmn/ J) are used. Mice are group housed at a constant temperature (23±1 °C) and in a 12-h light/12-h dark cycle (lights on 20:00–08:00). Adult DAT-Cre mice are anaesthetized by intraperitoneal injection of 1% Pentobarbital Sodium (60 mg/kg), then placed in a stereotaxic instrument (68025, RWD). After sterilized with 75% alcohol, incisions are made to expose the skull, align the skull to standard stereotaxic coordinates. A small hole with a diameter of ∼500 μm is formed by a drill on the skull, and then 500 nl virus (AAV2/9-hEF1a-DIO-hChR2(H134R)-mCherry-WPRE-pA) is injected by a glass micropipette using a microsyringe pump (Nanoliter 2000 injector with the Micro4 controller, WPI) at a rate of 46 nl/min into the VTA (stereotaxic coordinates from bregma (mm): anterior-posterior (AP): −3.40, medial-lateral (ML): +/− 0.50, dorsal-ventral (DV): −4.20) at rate of 46 nl/min. After suturing the wound, mice recover with the virus expressing for 3 weeks, a second surgery is operated to implant the three electrodes. The implantation procedure is similar to the injection procedure. After fixing the anaesthetized mouse in a stereotaxic instrument and exposing the skull, two holes are made by a drill. a stainless steel screw tied with a Ag wire is inserted into brain to work as the counter electrode. Dry the skull surface, and then the microprobe is implanted into VTA (stereotaxic coordinates from bregma (mm): anterior-posterior (AP): −3.40, medial-lateral (ML): +/− 0.52, dorsal-ventral (DV): −4.50). Dental cement is used to affix the probe and the stainless steel screw. A third hole is made on the skull after dental cement is dried, then a Ag/AgCl wire is implanted to work as the reference electrode, and dental cement is used again to affix the three electrodes. Both the CE and the RE are implanted elsewhere in the mouse brain cortex (Fig. S12). For the implantation of the probe used for RTPP, pictures of surgery procedure are provided (Fig. S13).

#### Immunohistochemistry

Mice are perfused by the phosphate buffered saline (PBS) solution and followed by 4% paraformaldehyde. Mouse brains are post-fixed in the same paraformaldehyde solution overnight and then immersed in 30% sucrose solution for 24 h. The frozen brains are cut into sections with a thickness of 40 μm, and then the sections are washed by PBS and PBST (0.3% Triton X-100 in PBS) and blocked in 3% bovine serum albumin (BSA) in PBST at room temperature for 1 h. The sections are incubated with primary antibodies (rabbit polyclonal Anti-Tyrosine Hydroxylase Antibody, 1:1000) for 24 h at 24 °C, then washed in PBST for 5 times and incubated with secondary antibodies (Goat anti-Rabbit IgG (H+L) Cross-Adsorbed Secondary Antibody, Alexa Fluor 488, 1:500) for 2 h at room temperature (avoid light), and finally washed the sections in PBS for 3 times. Fluorescent images are scanned with an automated slider scanner (17026728, Zeiss) and a laser scanning microscope (FV3000, Olympus).

## Acknowledgments

The research is supported by National Natural Science Foundation of China (NSFC) (61874064), Beijing Innovation Center for Future Chips, Tsinghua University, and Beijing National Research Center for Information Science and Technology (BNR2019ZS01005). X. S., M. F. and Q. S. are also grateful for the financial support from the CAS Interdisciplinary Innovation Team. We thank L. Sun and Z. Zhou (Peking University), L. Lu and M. Luo (National Institute of Biological Sciences) for valuable discussions. We also thank J. Wei and C. Li (University of Science and Technology Beijing) for the growth of diamond films.

## Author contributions

X. S. developed the concepts. C. L., Y. X., T. W., L. Li, R. L., Y. D., H. D., G. L., G. Z., L. Liu, G. Z., M. F., Q. S., L. Y., and X. S. performed device fabrication and characterization. C. L., C. D., Y. X., and X. S. performed simulations. Y. Z. and X. S. designed circuits. C. L., Y. Z., X. C., Y. X., and X. S. designed and performed biological experiments. C. L. and X. S. wrote the paper in consultation with all other authors.

## Competing interests

The authors declare that they have no competing interests.

## Data and materials availability

All data needed to evaluate the conclusions in the paper are present in the paper and/or the Supplementary Materials. Additional data related to this paper may be requested from the authors.

## Supplemental Information

### Device fabrication

The detailed structure of our proposed optoelectrochemical probe involves (from bottom to top): an flexible double side copper (Cu) coated polyimide (PI) (18 μm Cu / 25 μm PI / 18 μm Cu) substrate, an indium gallium nitride (InGaN) based blue emitting micro-LED (size: 125 μm × 185 μm × 7 μm), an undoped diamond interlayer (size: 180 μm × 240 μm × 20 μm) and a PEDOT:PSS film (size: 150 μm × 200 μm × 0.1 μm).

A detailed description of the process to fabricate the implantable, fully integrated microprobe system is listed below:

#### LED Fabrication

The LED structure (from bottom) included the sapphire substrate, a GaN buffer layer, an n-GaN, an InGaN/GaN multiple-quantum-well layer, and a p-GaN. The processes for LED fabrication are the same as those described our previous work.

Reference: Li, L. et al. Heterogeneous Integration of Microscale GaN Light-Emitting Diodes and Their Electrical, Optical, and Thermal Characteristics on Flexible Substrates. *Advanced Materials Technologies* **3**, 1700239 (2018).

#### Preparation of the Adhesive Solution

The adhesive solution comprises a mixture of:

bisphenol A glycerolate (1 glycerol/phenol) diacrylate;
3-(Trimethoxysilyl) propyl methacrylate;
Spin-on-Glass (SOG 500F, Filmtronics Inc.);
2-Benzyl-2-(dimethylamino)-4’-morpholinobutyrophenone;
and anhydrous ethanol.
The weight ratio is 200:100:100:9:2000. Stir at room temperature until full mixing.
Store the mixed solution in refrigerator (4 °C) for future use.

Reference: Kim, T. et al. Thin Film Receiver Materials for Deterministic Assembly by Transfer Printing. *Chemstry of Materials* **26**, 3502 (2014).

#### Diamond Fabrication

1. The undoped diamond film is grown on a silicon substrate by chemical vapor deposition (CVD).
2. Clean the grown diamond with acetone, isopropyl alcohol (IPA), deinoized (DI) water.
3. Laser cutting (Nd:YVO_4_ laser, 1064 nm) with a power of 7 W, pulse repetition rate of 1 MHz, scan speed of 1 m/s and repeat scan for 200 times.
4. Etch the Si substrate in CH_3_COOH : HNO_3_ : HF = 5:5:2 by volume for 0.5 hour to release the diamond film, and rinse with DI water.

#### PEDOT:PSS Preparation

5. Clean a brown glass bottle with standard RCA clean process 1 (NH_4_OH : H_2_O_2_ : H_2_O = 1:1:5, 80 °C) for 10 min.
6. Fully mix 10 ml 100.00% PEDOT:PSS (CLEVIOS PH 1000, Xi’an Polymer Light Technology Corp.) with 0.5 ml of Ethylene glycol (AR, Beijing Lanyi chemical products Co., Ltd), 10μL of Dodecylbenzene sulfonate (DBSA) (90%, Shanghai Aladdin Bio-Chem Technology Co., LTD) and 0.1ml 3-Glycidoxypropyltrimethoxysilane (GOPS) (97%, J&K Scientific Ltd.) for 1 h.
7. Store the mixed solution in refrigerator (4 °C) for future use.

#### LED transfer printing

8. Clean a glass substrate with standard RCA clean process 1 for 10 min.
9. Dehydrate at 110 °C for 10 min.
10. Spin-coat with poly(dimethylsiloxane) (PDMS, Sylgard 184, pre-polymer : curing agent = 10:1, by weight, 500 rpm / 6 s, 3000 rpm / 30 s) and soft-bake at 110 °C for 25 seconds.
11. Laminate a flexible double side copper (Cu) coated polyimide (PI) (18 μm Cu / 25 μm PI / 18 μm Cu, DuPont) substrate on the glass and post-bake at 110 °C for 10 min.
12. Clean the double side copper (Cu) coated polyimide (PI) film with acetone, isopropyl alcohol (IPA), deinoized (DI) water and dehydrate at 110 °C for 10 min.
13. Spin coat 10 µm thick polyimide (YDPI-102, YiDun New Material Suzhou), baked at 250 °C for more than 2 hours.
14. Spin coat negative photoresist (PR) (AZ nLOF 2070, 500 rpm / 5 s, 3000 rpm / 30 s) and soft-bake at 110 °C for 2 min.
15. Expose with 365 nm optical lithography with irradiance for 45 mJ/cm^2^ (URE-2000/25, IOE CAS) through a chrome mask and post-exposure bake at 110 °C for 90s.
16. Develop PR in aqueous base developer (AZ300 MIF) and rinse with DI water.
17. Deposit 10 nm / 100 nm of Cr/Al as the markers by sputter coater.
18. Lift-off PR in acetone.
19. Clean the processed substrate in step 18 (acetone, IPA, DI water).
20. Dehydrate at 110 °C for 10 min.
21. Spin-coat with adhesive liquid (3000 rpm, 30 s) on the substrate and soft-bake at 110 °C for 4 min.
22. Transfer printing the LED from the source wafer onto the processed substrate with PDMS stamp.
23. Cure under to ultraviolet (UV) for 1 h and bake at 110 °C for 1 h.

#### Epoxy encapsulation (LED)

24. Clean the processed wafer in step 23 (acetone, IPA, DI water) and dehydrate at 110 °C for 10 min.
25. Expose to ultraviolet induced ozone (UV Ozone) for 10 min.
26. Spin-coat with epoxy SU8-2002 (500 rpm/ 5 s, 3000 rpm/ 30 s).
27. Soft-bake at 65 °C for 1 min and 95 °C for 1 min.
28. Pattern epoxy to reveal contact pads of LED by exposing with UV lithography with irradiance for 100 mJ/cm^2^ through a chrome mask.
29. Post-bake at 65 °C for 1 min and 95 °C for 2 min.
30. Develop in propylene glycol monomethyl ether acetate (PGMEA) for 1 min and rinse with IPA.
31. Hard bake at 110 °C for 20 min.

#### LED interconnect metallization

32. Clean the processed wafer in step 31 (acetone, IPA, DI water) and dehydrate at 110 °C for 10 min.
33. Pattern PR AZ nLOF 2070.
34. Deposit 10 nm / 600 nm / 200 nm of Cr/Cu/Au by sputter coater.
35. Lift-off PR in acetone.

#### LED encapsulation

36. Clean the processed wafer in step 35 (acetone, IPA, DI water) and dehydrate at 110 °C for 10 min.
37. Expose to ultraviolet induced ozone (UV Ozone) for 10 min.
38. Spin-coat with epoxy SU8-3005 (500 rpm/ 5 s, 3000 rpm/ 30 s).
39. Soft-bake at 65 °C for 1 min and 95 °C for 2 min.
40. Pattern epoxy by exposing with UV lithography tools with irradiance for 150 mJ/cm^2^ through a chrome mask.
41. Post-bake at 65 °C for 1 min and 95 °C for 3 min.
42. Develop in propylene glycol monomethyl ether acetate (PGMEA) for 1 min and rinse with IPA.
43. Hard-bake at 110 °C for 20 min.

#### Diamond transfer printing

44. Clean the processed wafer in step 43 (acetone, IPA, DI water) and dehydrate at 110 °C for 10 min.
45. Spin-coat with SU8-2002 (500 rpm/ 5 s, 3000 rpm/ 30 s).
46. Transfer printing the diamond from the source wafer onto the processed substrate with PDMS stamp.
47. Cure under to UV for 30 min and bake at 110 °C for 30 min.

#### PEDOT:PSS interconnect metallization

48. Clean the processed wafer in step 47 (acetone, IPA, DI water) and dehydrate at 110 °C for 10 min.
49. Spin-coat with SU8-3005 (500 rpm/ 5 s, 3000 rpm/ 30 s) and soft-bake at 65 °C for 1 min and 95 °C for 2 min.
50. Thicken the SU8-3005 by a second spin-coat process.
51. Pattern PR SU8-3005 (expose diamond) to buffer the height difference caused by diamond thickness, and hard-bake at 110 °C for 20min.
52. Clean the processed wafer in step 51 (acetone, IPA, DI water) and dehydrate at 110 °C for 10 min.
53. Pattern PR AZ nLOF 2070.
54. Roughen surface by reactive ion etching (RIE) with oxygen gas (O_2_, 100 sccm, 90 mTorr, 150 W) for 30s.
55. Deposit 500 nm of Au by sputter coater.
56. Lift-off PR in acetone.

#### PEDOT:PSS coating

57. Clean the processed wafer in step 56 (acetone, IPA, DI water) and dehydrate at 110 °C for 10 min.
58. Expose to ultraviolet induced ozone (UV Ozone) for 10 min.
59. Spin-coat with PEDOT:PSS (500 rpm/ 5 s, 2000 rpm/ 30 s) and bake at 110 °C for 1 h.
60. Clean the processed wafer in step 59 (acetone, IPA, DI water) and dehydrate at 110 °C for 10 min.
61. Spin coat positive PR (SPR220-3.0, Microchem, 500 rpm / 5 s, 3000 rpm / 30 s) and soft-bake at 110 °C for 1.5 min.
62. Pattern PR SPR220-3.0 by exposing with UV lithography tools with irradiance for 300 mJ/cm^2^ through a chrome mask and post-bake at 110 °C for 1.5 min.
63. Develop PR in aqueous base developer (AZ300 MIF), rinse with DI water and hard-bake at 110 °C for 10 min.
64. Clean PEDOT:PSS without PR covering by RIE with oxygen gas (O_2_ 100 sccm, SF_6_ 5 sccm, 90 mTorr, 150 W) for 2 min.
65. Lift-off PR in acetone.

#### Encapsulation and laser milling

66. Clean the processed wafer in step 65 (acetone, IPA, DI water) and dehydrate at 110 °C for 10 min.
67. Expose to ultraviolet induced ozone (UVO) for 10 min.
68. Pattern epoxy SU8 3005 (500 rpm/ 5 s, 3000 rpm/ 30 s), cure under to UV for 150 mJ/cm^2^ and bake at 100 °C for 30 min.
69. A bilayer of SU8 3005 by a second spin-coat process.
70. UV laser milling to form the probe shape, release from the glass substrates.

**Figure S1.**
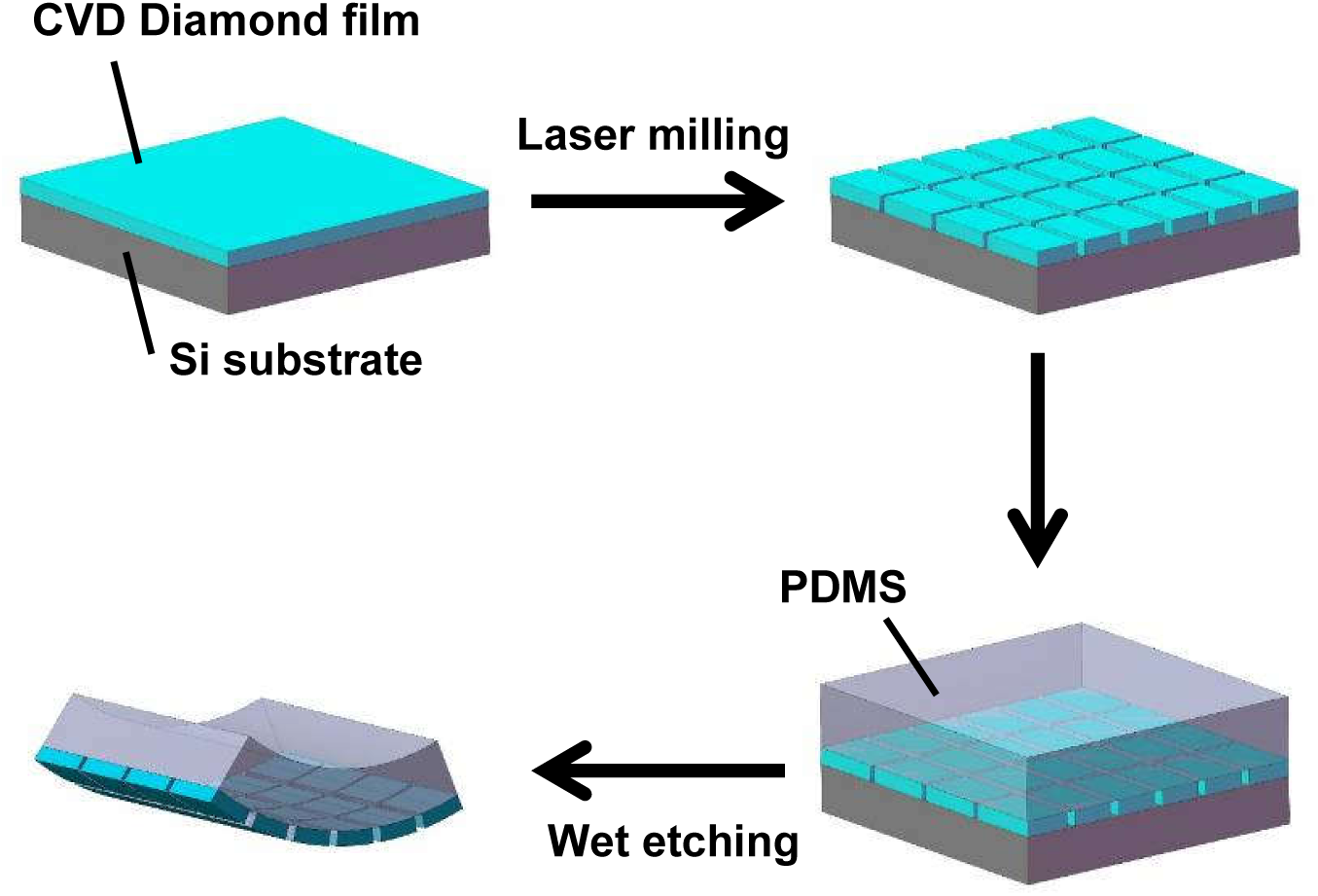
Schematic illustration of processing flow for diamond film fabrication, including chemical vapor deposition, laser milling, Si subtrate etching and transfer printing.

**Figure S2.**
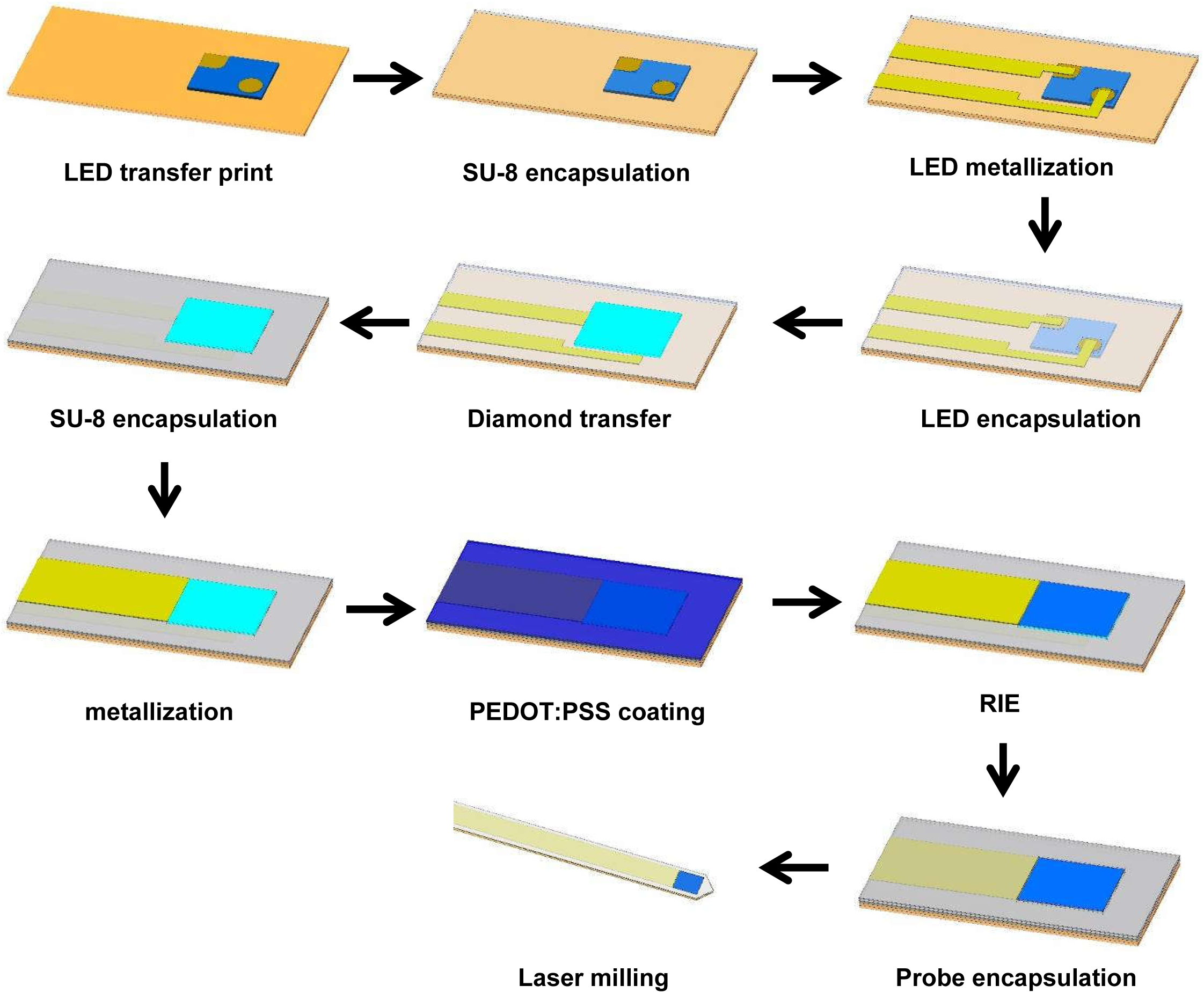
Schematic illustration of processing flow for the optoelectrochemical probe fabrication.

**Figure S3.**
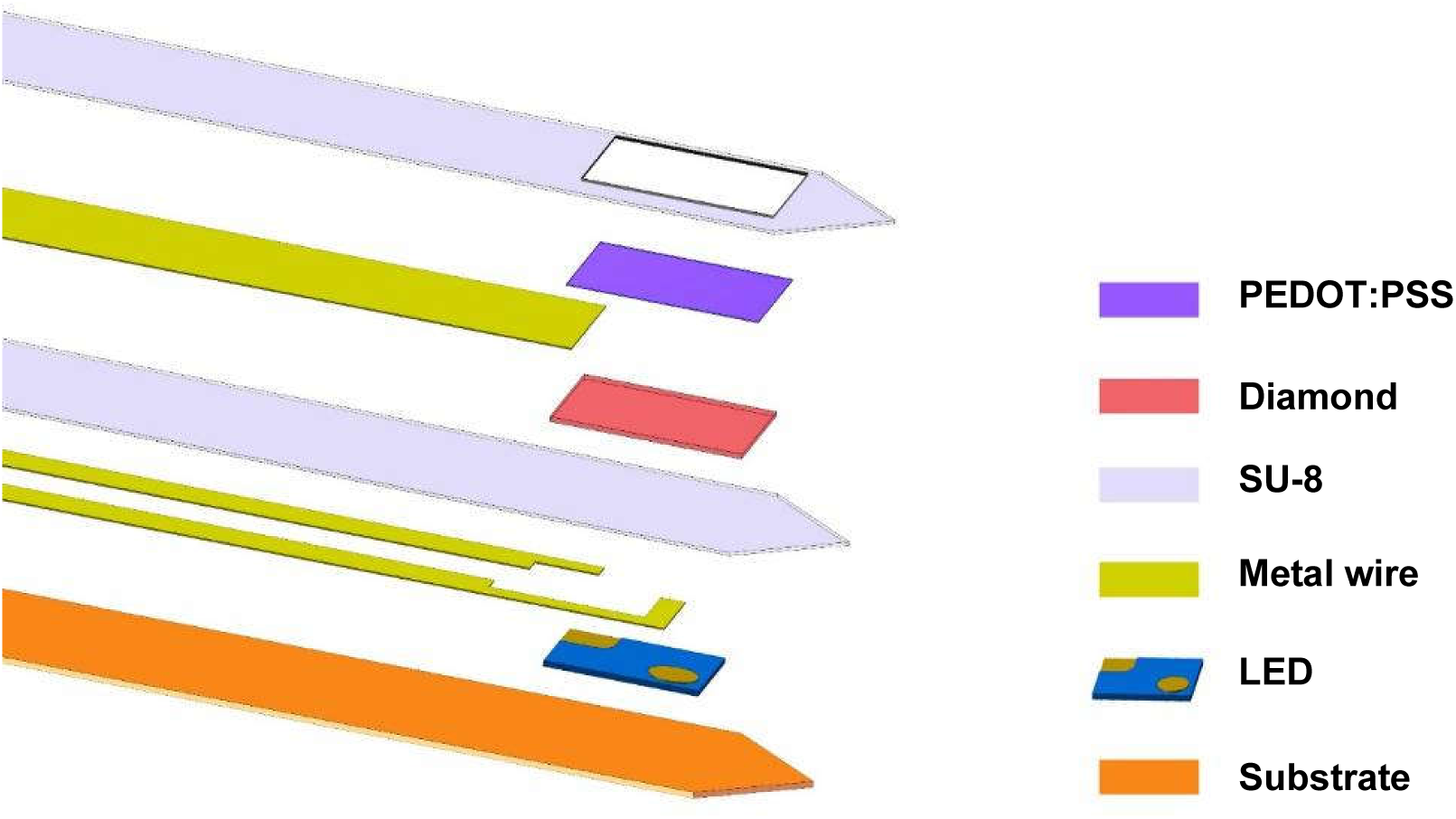
Exploded schematic illustration of the microprobe, including the details of the metallized wires and SU-8 encapsulation layers.

**Figure S4.**
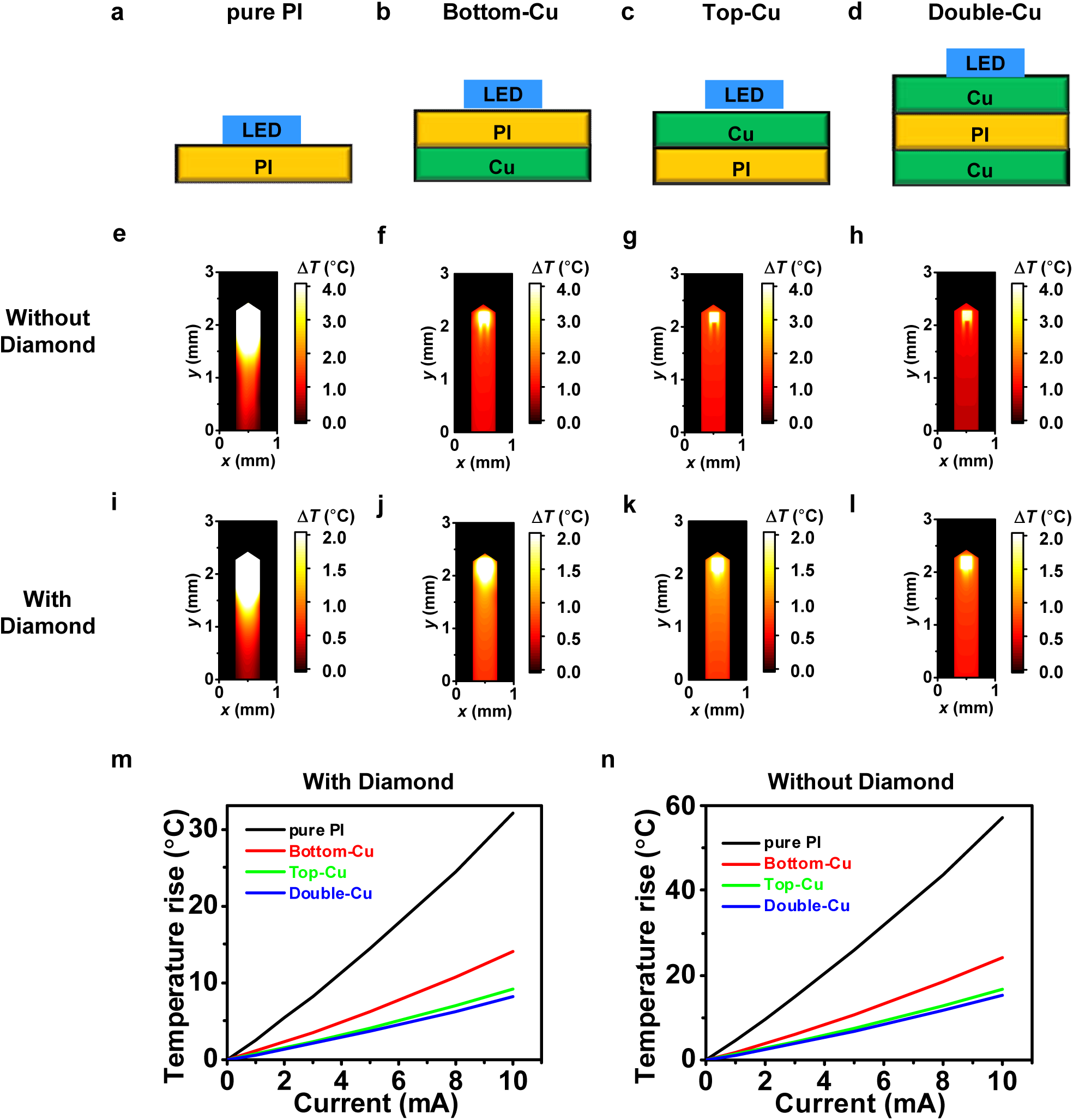
The schematic illustration of a micro-LED on **(a)** a pure polyimide (PI) substrate, **(b)** bottom Cu coated PI, **(c)** top Cu coated PI, and **(d)** double side Cu coated PI. **(e-l)** Simulation of temperature distributions of the LED probes operated in air (current 3 mA, frequency 20 Hz and duty cycle 20%) based on various substrates with and without diamond/PEDOT:PSS coatings. **(m, n)** Simulated maximum temperature rises above room temperature on the top surface of the LED probes based on different substrates as a function of pulsed currents (frequency 20 Hz and duty cycle 20%) (m) with and (n) without diamond/PEDOT:PSS coatings.

**Figure S5.**
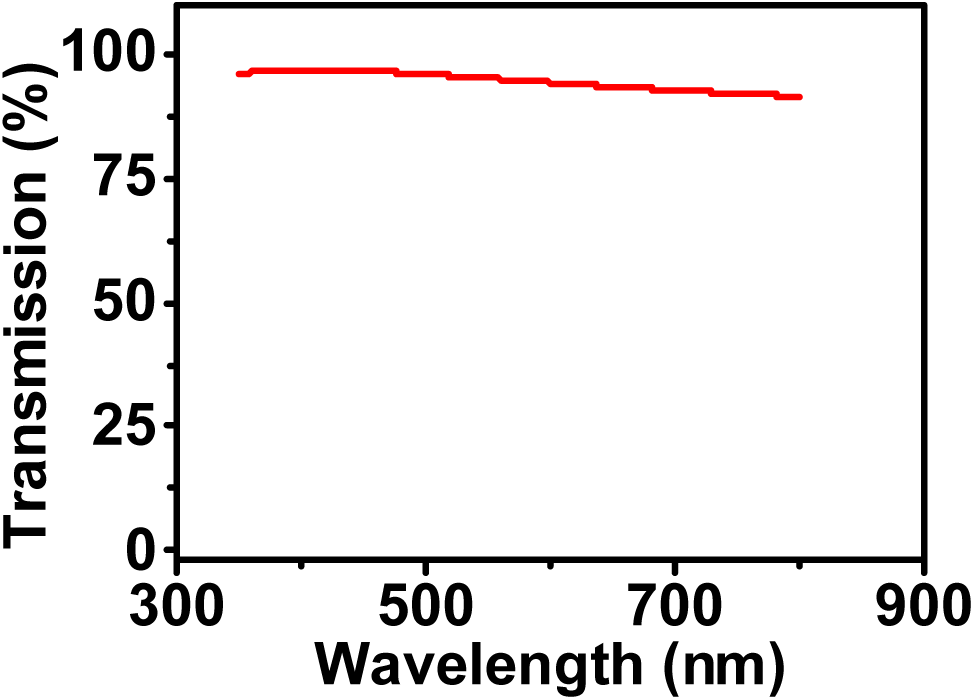
Optical transmission spectrum of a PEDOT:PSS film (about 100 nm thick) coated on glass.

**Figure S6.**
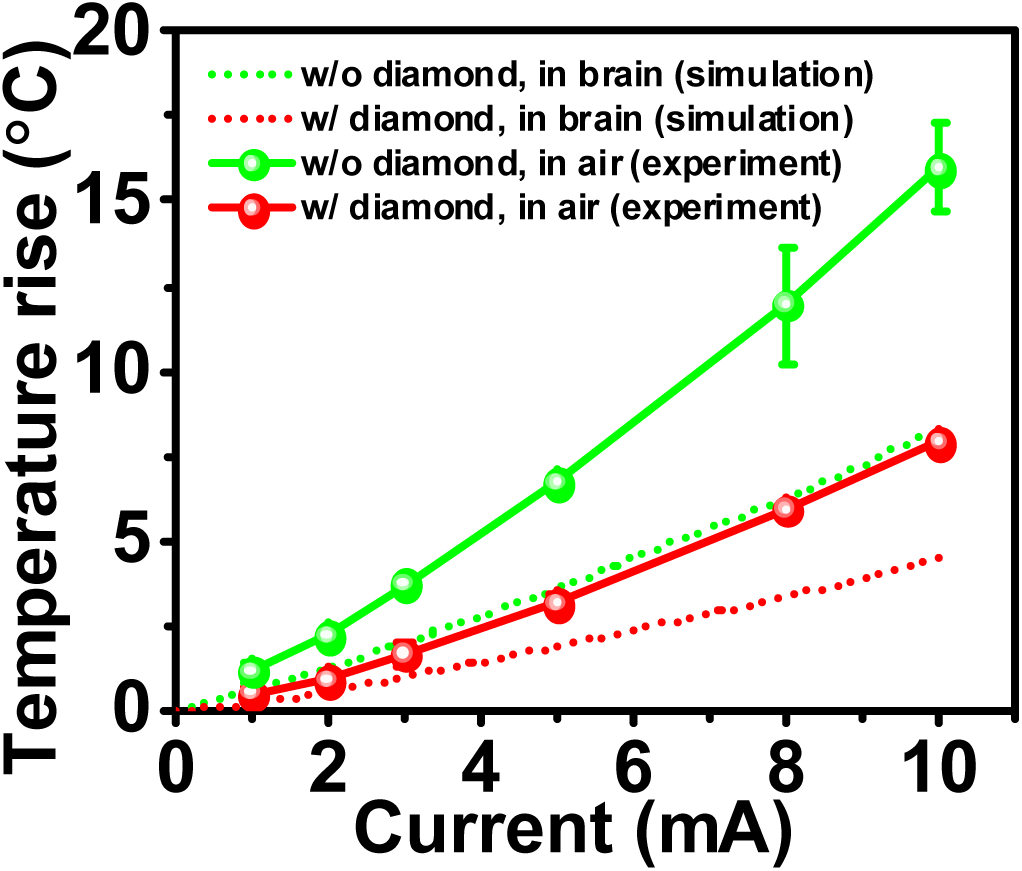
The predicted temperature rise of probe in brain tissue as a function of pulsed currents (frequency 20 Hz and duty cycle 20%) based on finite element analysis. Solid line is experiment data obtained in air as shown in Fig. 2d, dot line is the simulation data; green is without diamond, red is with diamond.

**Figure S7.**
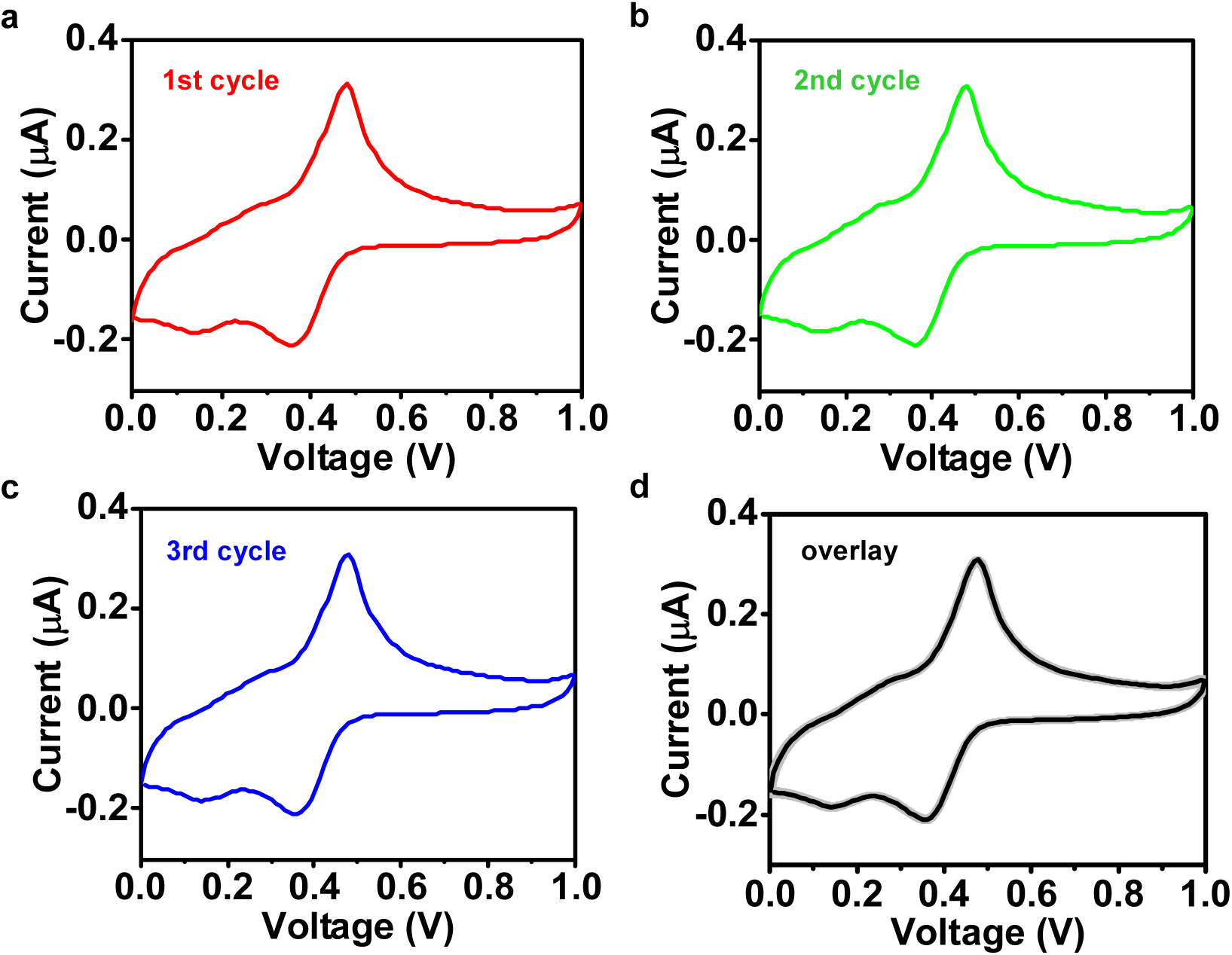
(a-c) Repeated CV scanning curves for a representative PEDOT:PSS probe (10 μM dopamine in HCl solution, pH = 4.0). **(d)** Overlay of the three curves.

**Figure S8.**
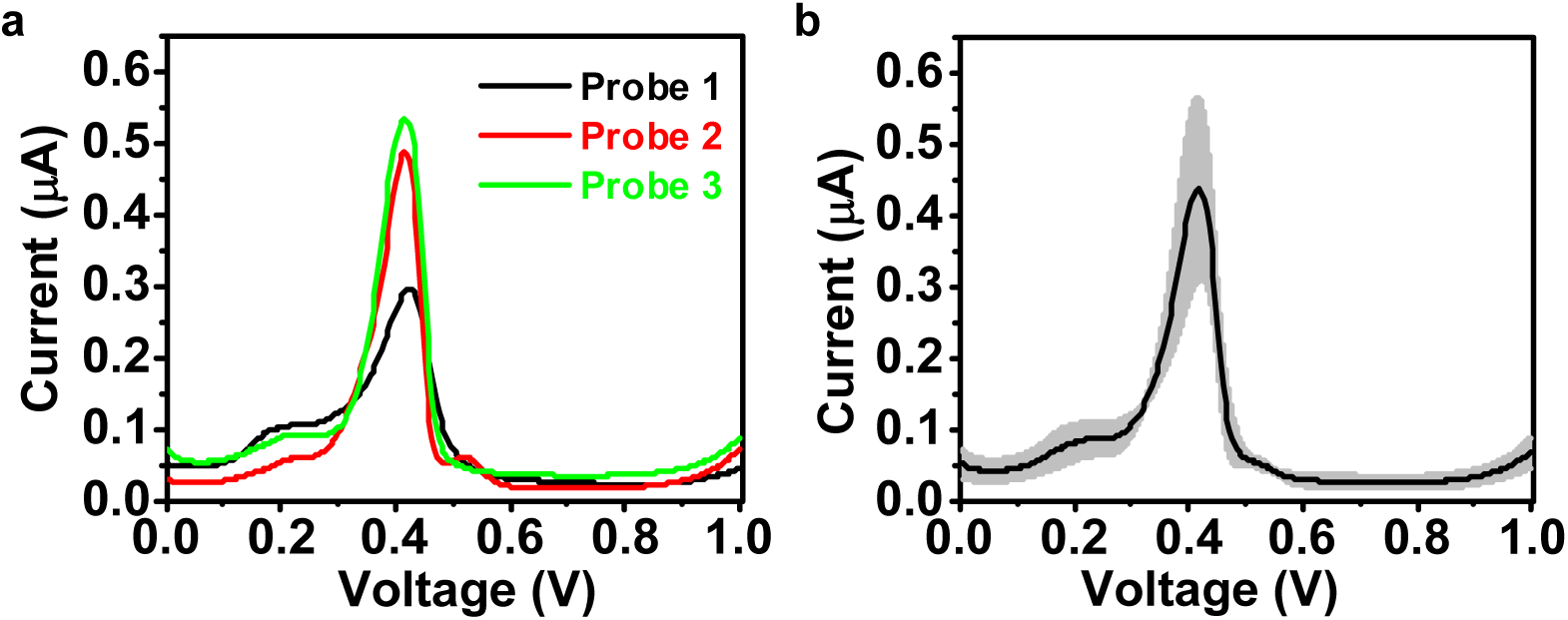
(a) DPV curves for 3 PEDOT:PSS probes (10 μM dopamine in HCl solution, pH = 4.0). **(b)** average of the 3 curves.

**Figure S9.**
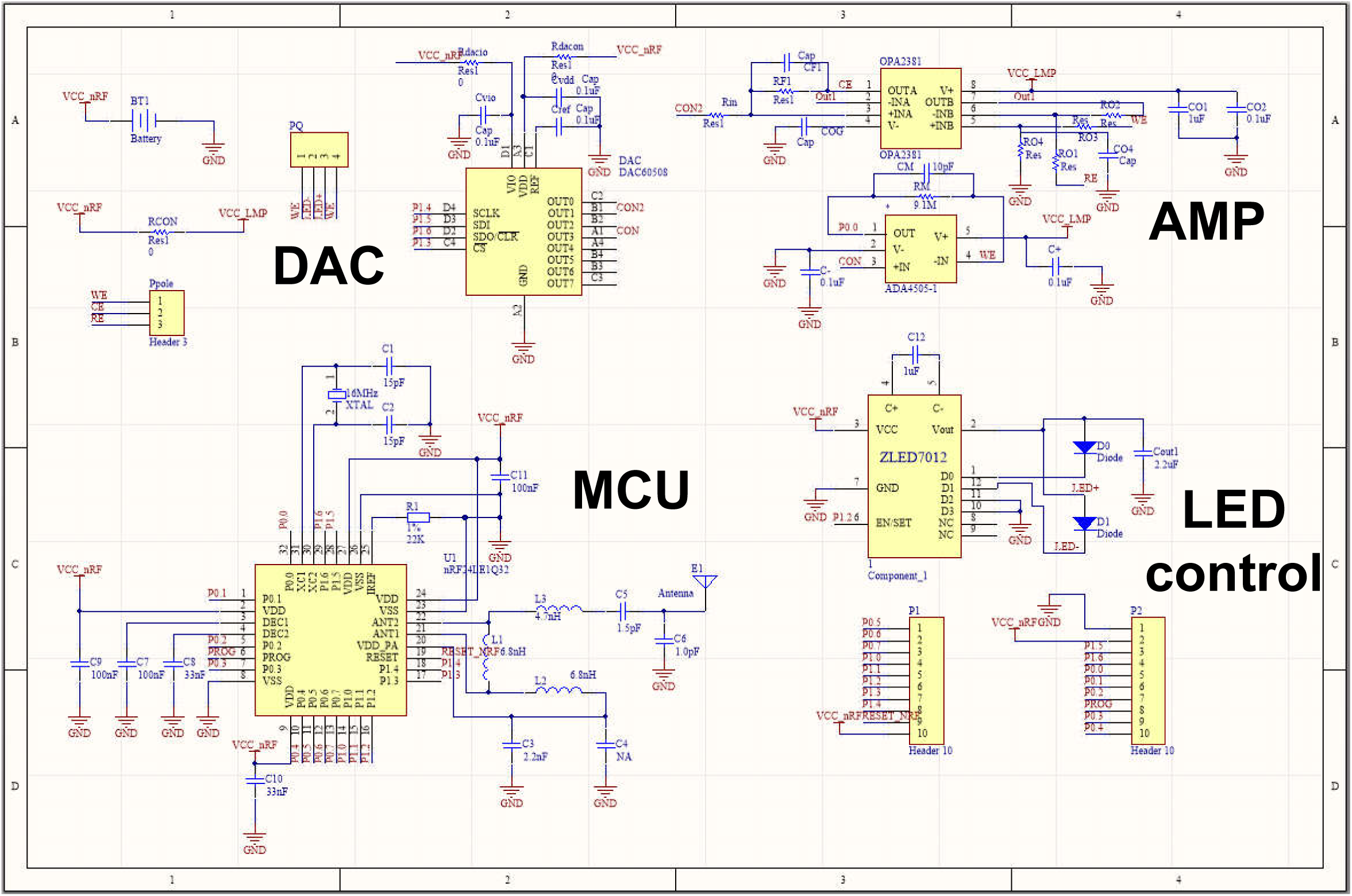
Schematic diagram of the wireless control circuit for the microprobe.

**Figure S10.**
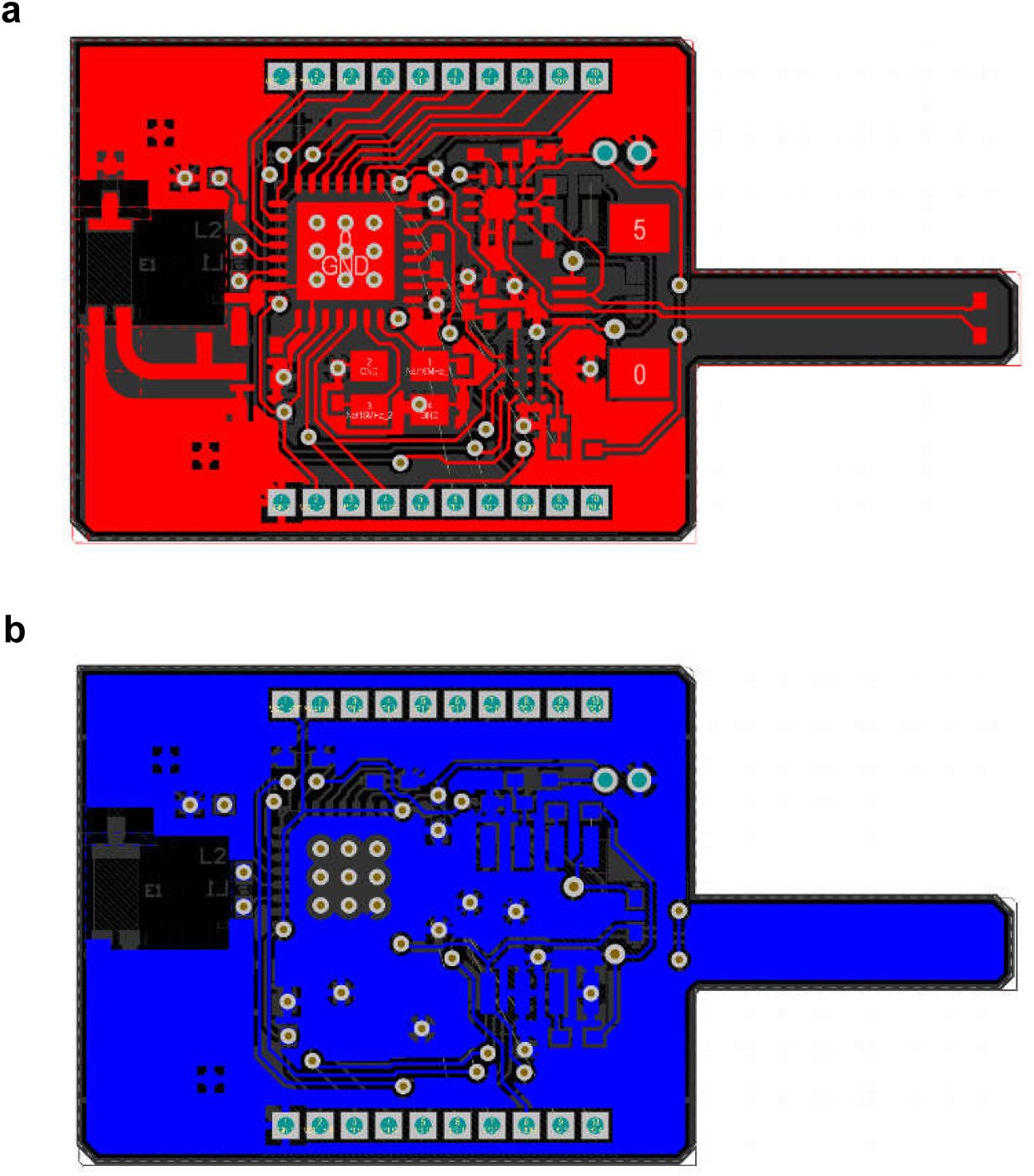
Layout of the designed printed circuit board. a) front view and b) back view.

**Figure S11.**
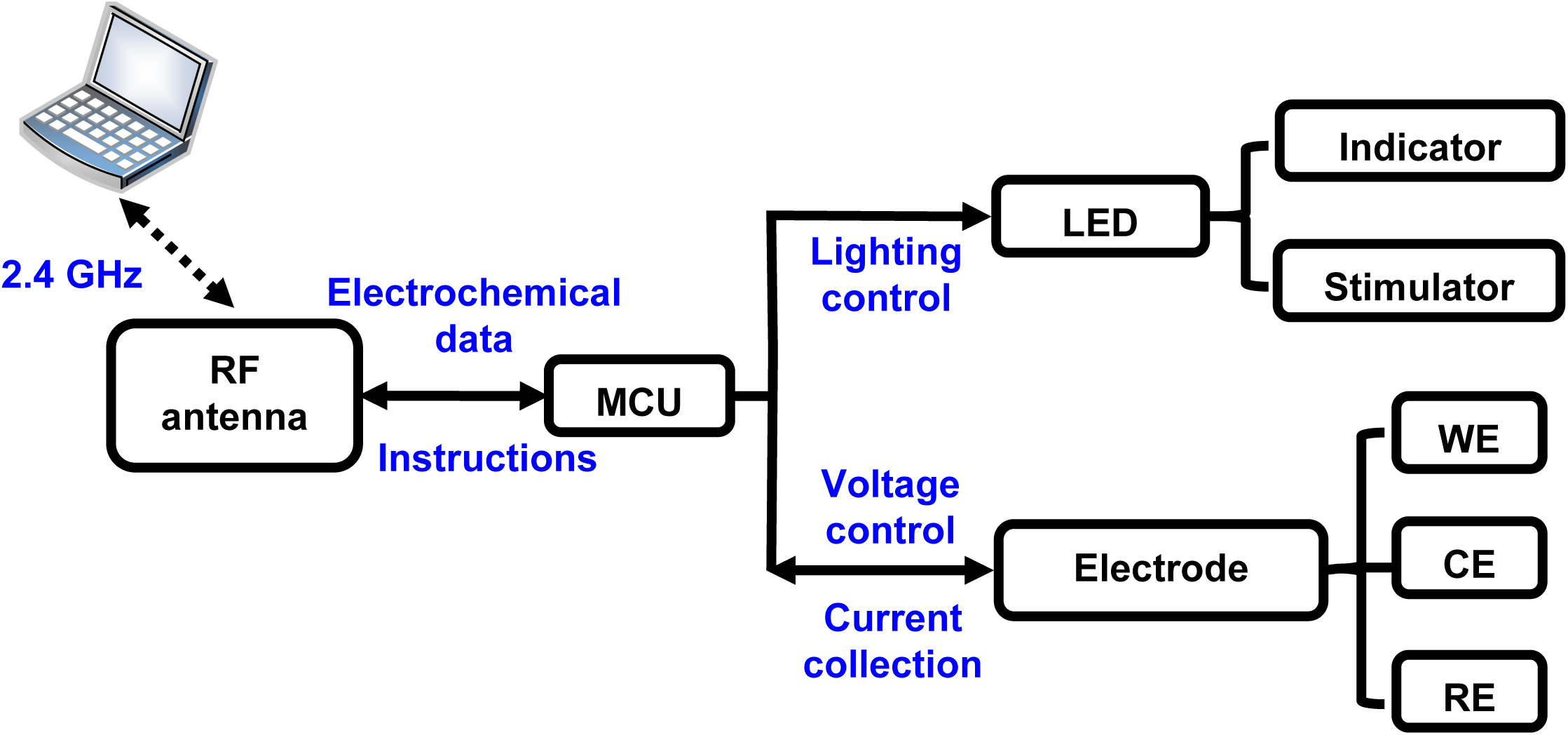
Schematic diagram of the wireless system (WE: working electrode, CE: counter electrode, RE: reference electrode).

**Figure S12.**
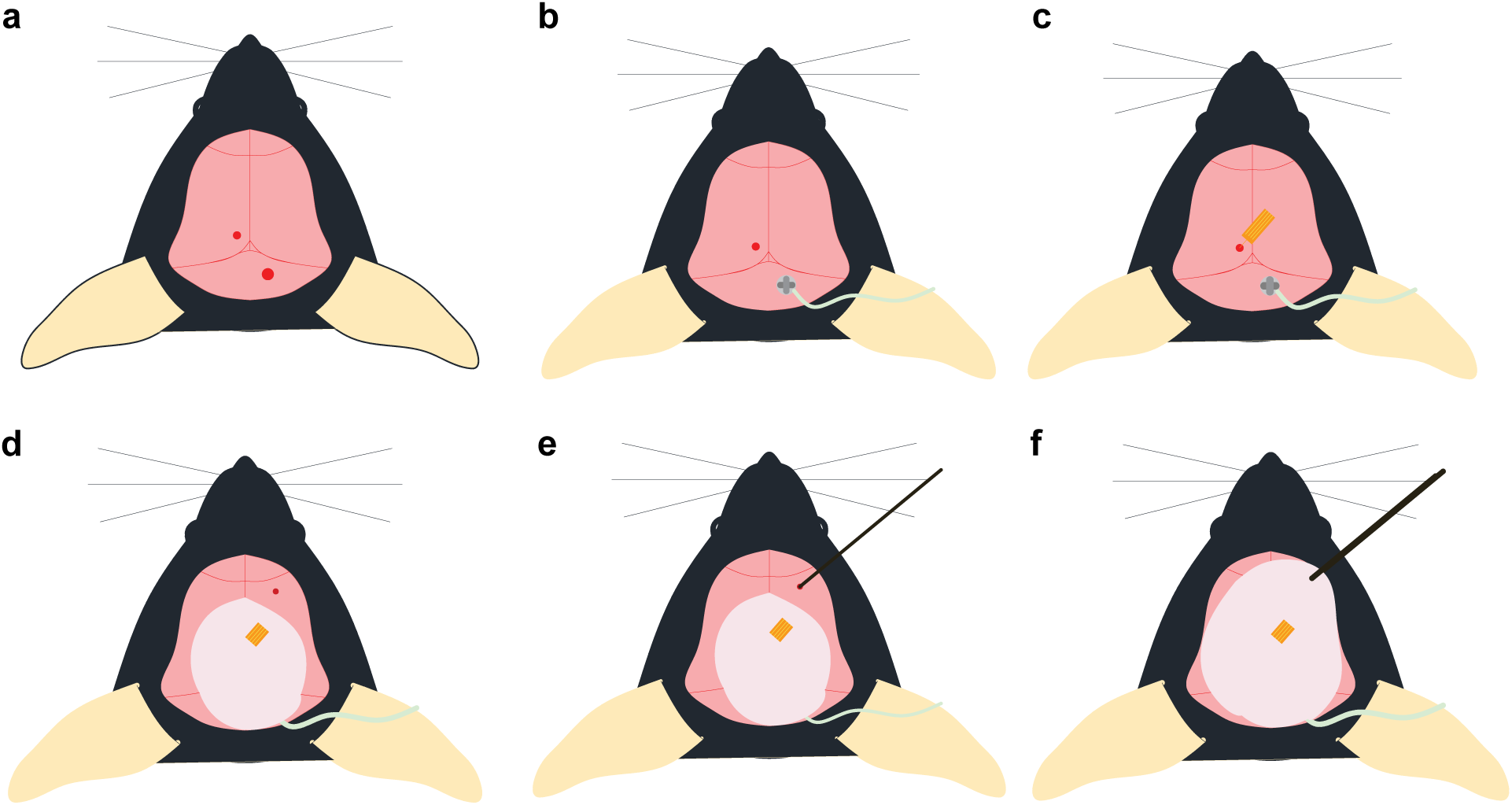
Schematic diagram of experimental procedure of implanting the three electrodes. a) exposing the mouse skull and two holes are made by a drill; b) inserting a stainless steel screw tied with a Ag wire into brain; c) implanting the microprobe into VTA; d) dental cement is used to affix the probes and the stainless steel screw, and third hole is made on the skull; e) implanting a Ag/AgCl wire to works as the reference electrode; f) more dental cement is used to affix the three electrodes.

**Figure S13.**
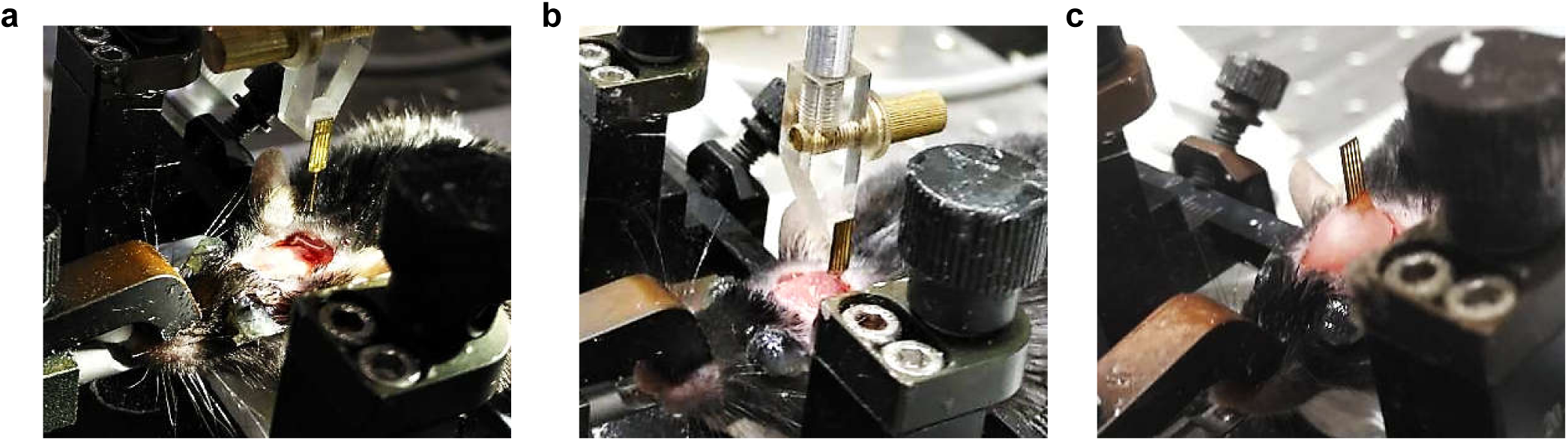
Experimental procedure of implanting the flexible probe. a) expose the mouse skull and a small hole is made by a drill; b) implant the microprobe into VTA, and apply dental cement fix the probe; c) release the holder after dental cement is dry.

**Figure S14.**
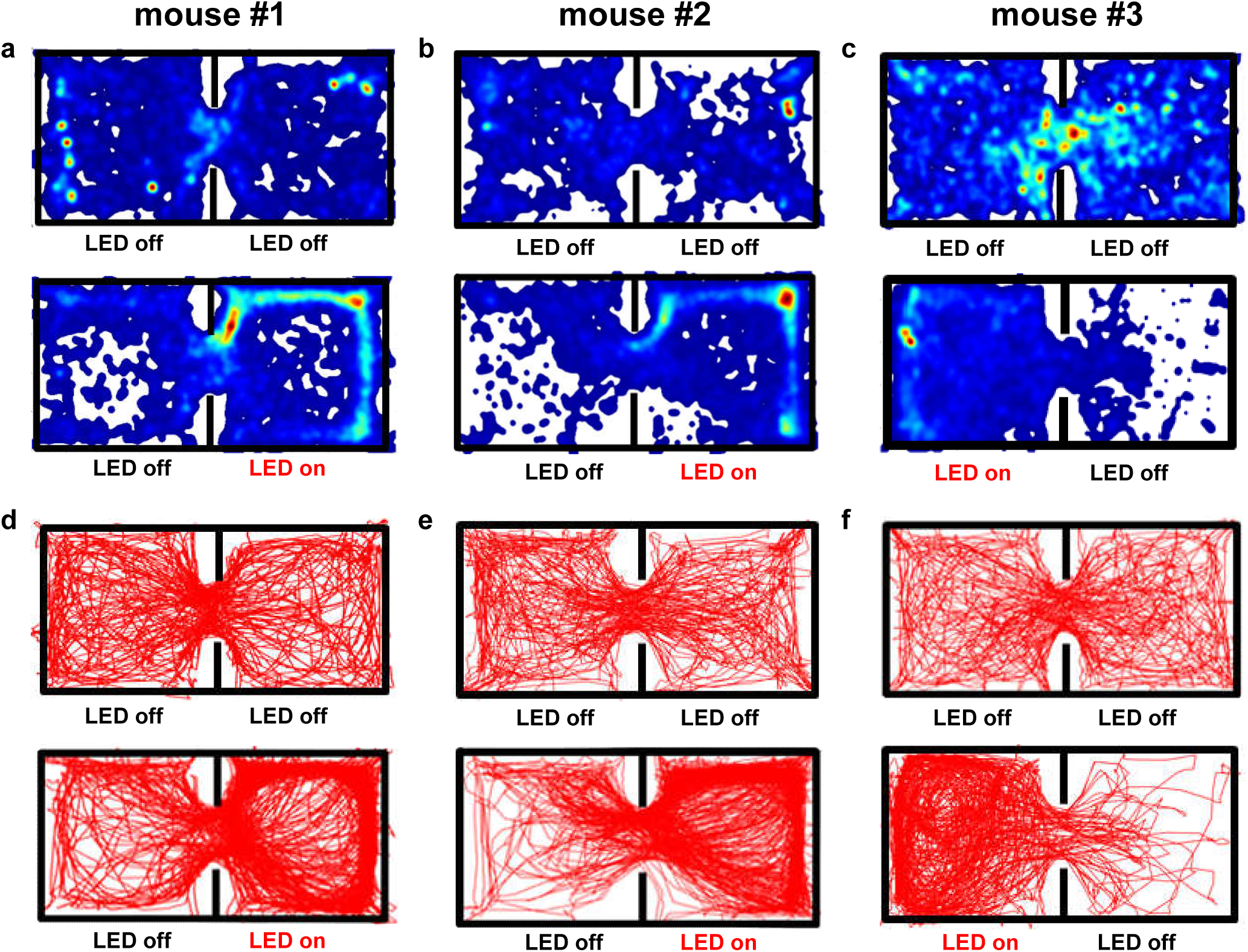
Position heat map (a–c) and corresponding moving traces (d–f) for three different mice, during pretests and under optogenetic stimulations.

**Figure S15.**
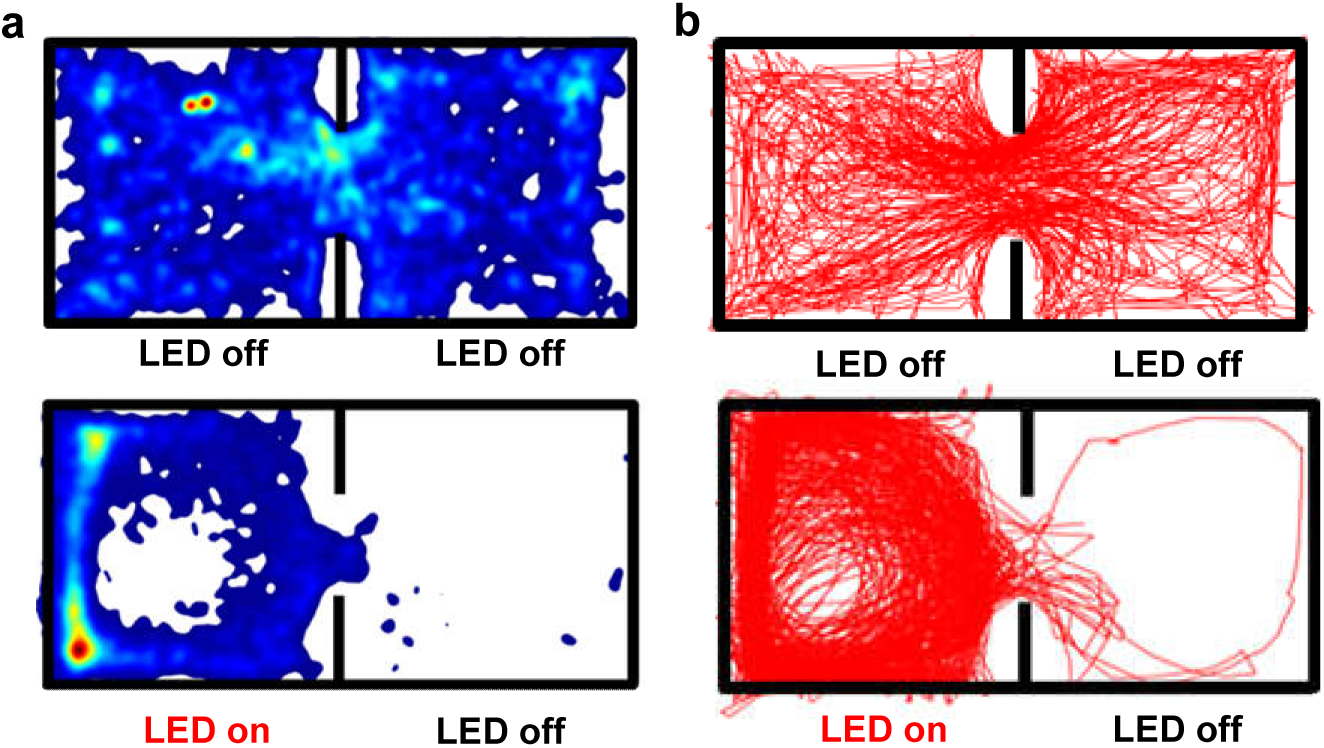
a) position heat maps and moving traces for mouse #2 in Fig. S14, during pretests and under optogenetic stimulations, which are recorded 17 days after implanting probe. The position pereference is more evident than results in Fig. S14, because of the residual reward memory.

**Figure S16.**
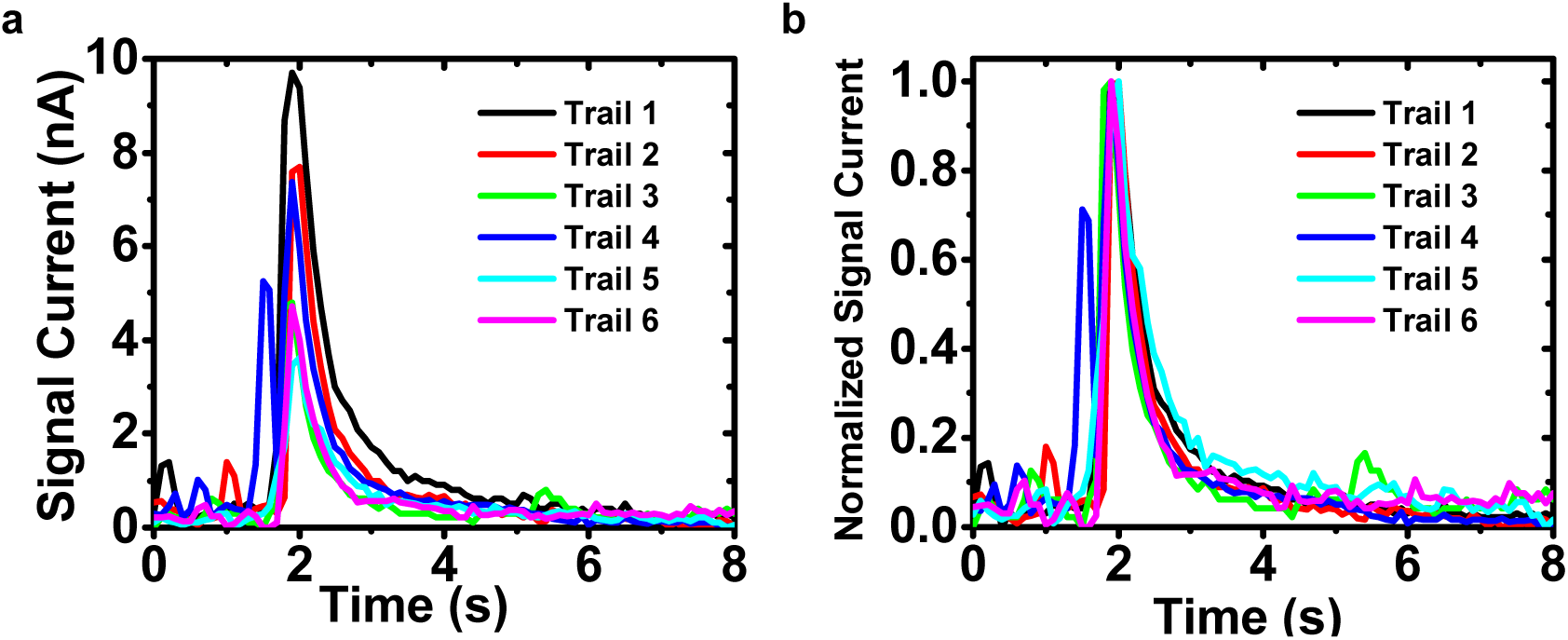
*In Vivo* electrochemical response signals (6 representative trails) collected from 2 mice. **(a)** Current signals obtained by subtracting baseline currents. **(b)** Normalized current signals.

**Figure S17.**
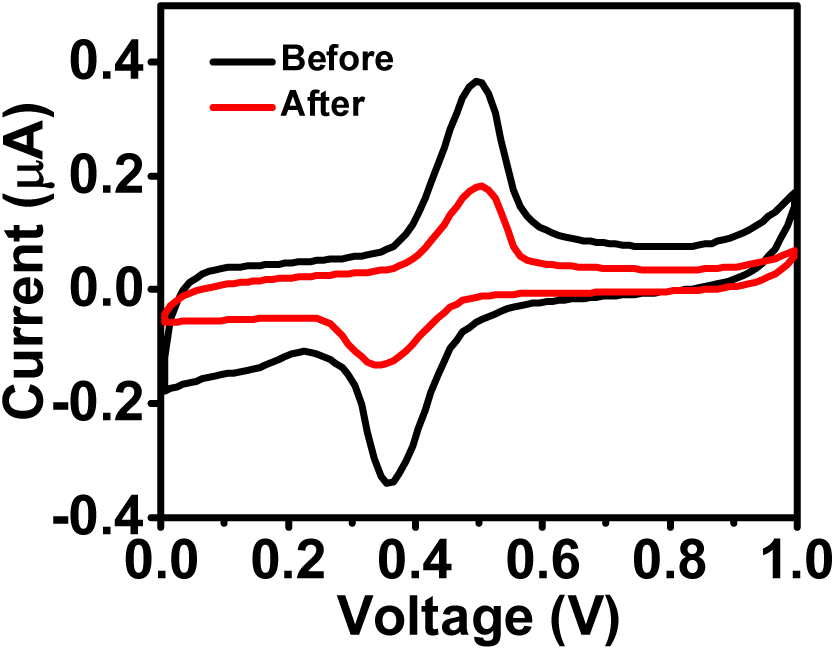
CV scanning curves for a representative probe (1 mM dopamine in HCl solution, pH = 4.0) before and 1 day after the implantation into the mouse brain for in vivo tests, showing the performance degradation in the living animals.

**Figure S18.**
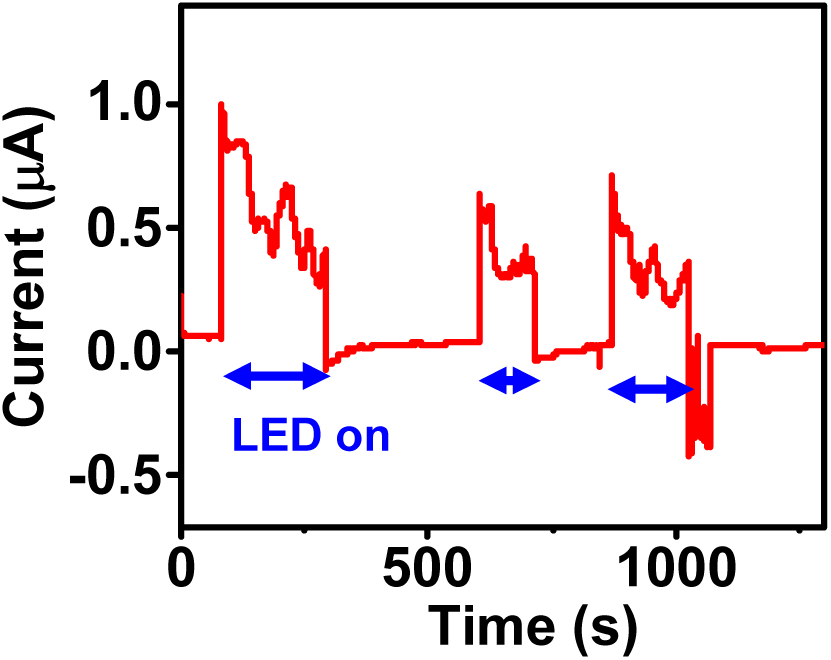
Interference of the recorded electrochemical signals (CA test *in vitro*) by the optical stimulation.

## Notes

### Competing Interest Statement

The authors have declared no competing interest.

